# Hybrid lipid-block copolymer membranes enable stable reconstitution of a wide range of nanopores and robust sampling of serum

**DOI:** 10.1101/2024.05.16.594548

**Authors:** Edo Vreeker, Fabian Grünewald, Nieck Jordy van der Heide, Siewert-Jan Marrink, Katarzyna (Kasia) Tych, Giovanni Maglia

## Abstract

Biological nanopores are powerful tools for detecting biomolecules at the single-molecule level, making them appealing as sensors for biological samples. However, the lipid membranes in which nanopores reside can be unstable in the presence of biological fluids. Here, membranes formed with the amphiphilic polymers PMOXA-PDMS-PMOXA and PBD-PEO are tested as potential alternatives for nanopore sensing. We demonstrate that polymer membranes can possess increased stability towards applied potentials and high concentrations of human serum, but that the stable insertion of a wide range of biological nanopores is most often compromised. Alternatively, hybrid polymer-lipid membranes comprising a 1:1 w/w mixture of PBD_11_PEO_8_ and DPhPC showed high electrical and biochemical stability while creating a suitable environment for all tested nanopores. Analytes such as proteins, DNA and sugars were efficiently sampled, indicating that in hybrid membranes nanopores showed native-like properties. Molecular dynamics simulations revealed that lipids form ∼12 nm domains interspersed by a polymer matrix. Nanopores partitioned into these lipid nanodomains and sequestered lipids possibly offering the same binding strength as in a native bilayer. This work shows that single-molecule analysis with nanopores in [PBD_11_PEO_8_ + DPhPC] membranes is feasible and present stable recordings in the presence of human serum. These results pave the way towards novel nanopore-based biosensors.

## Introduction

Biological nanopores have emerged as reliable tools for detecting metabolites and biomarkers directly from biological fluids.^[1–6]^ For this purpose, nanopores are typically reconstituted in planar membranes formed from DPhPC lipids (1,2-diphytanoyl-sn-glycero-3-phosphocoline). ^[7,8]^ However, it is well-known that biological fluids like human serum contain various components that interact with the lipid bilayer, often resulting in membrane rupture. ^[9,10]^ As a consequence, nanopore recordings of such samples are limited to the short lifetime of the membrane, creating a bottleneck for the development of novel nanopore biosensors. It is therefore of interest to develop a more robust membrane that enables stable nanopore recording in such challenging conditions.

Amphiphilic block copolymers can form substitutes for lipid systems and have been widely studied as novel nanocarriers and drug delivery vehicles.^[11,12]^ Two commercially available amphiphilic block copolymers are the triblock polymer PMOXA-PDMS-PMOXA (poly(2-methyloxazoline-b-dimethylsiloxane-b-2-methyloxazoline)) and diblock polymer PBD-PEO (poly(1,2-butadiene)-b-poly(ethylene oxide)). Like lipids, these amphiphiles can self-assemble into a rich variety of macromolecular structures, such as micelles, monolayers, bilayers, and polymersomes.^[13–16]^ Importantly, these polymeric assemblies can show increased stability in biological environments compared to lipid aggregates.^[17–20]^

Various studies have been carried out in which membrane proteins are reconstituted into PMOXA-PDMS-PMOXA or PBD-PEO membranes, but information about the electrical properties of nanopores in these membrane environments is scarce.^[21–28]^ Only for two nanopores with β-barrel transmembrane regions, α-hemolysin (α-HL) and mycobacterium smegmatis porin A (MspA), electrophysiology data in both polymer membranes are reported. However, nanopores come in different sizes and shapes and can have α-helical or β-sheet transmembrane domains, meaning that insights obtained with α-HL and MspA may not be transferable to other nanopores. Moreover, single-molecule nanopore measurements in polymer membranes have only been carried out with DNA. ^[24–26]^

Importantly, whether the reconstitution of nanopores in polymer membranes enables stable detection of analytes from biological samples has not yet been explored. If the increased biochemical stability of the polymer membranes indeed allows nanopore recordings with biofluids for substantially longer times than lipid membranes, this may overcome an important bottleneck and aid in the development of novel nanopore-based biosensors.

In this work we investigate the compatibility of seven nanopores with both α-helical and β-sheet transmembrane domains in PMOXA-PDMS-PMOXA and PBD-PEO planar membranes and test how such membranes react to high applied electric potentials and exposure to high concentrations of human serum. We found that polymer membranes can have a strongly improved tolerance towards high applied potentials and high serum concentrations, but that these membranes were often not compatible with nanopores. Importantly, however, hybrid [PBD-PEO + DPhPC] membranes showed the best of both worlds, providing high electrical and biochemical stability coupled to low-noise electrical recordings with nanopores. Atomistic and coarse-grained molecular dynamics simulations revealed that reconstituted nanopores resided in DPhPC nanodomains interspersed by a polymer matrix. This membrane environment created a near-native lipid environment for nanopores to operate, while the interface benefited from stability properties provided by the block copolymers. Nanopores in hybrid membranes could distinguish a wide range of analytes and were stable when exposed to human serum at concentrations greatly exceeding the limit for lipid membranes, highlighting the potential of these hybrid membranes for novel nanopore-based biosensors.

## Results and Discussion

### Polymer membrane stability

In this study, planar membranes were formed according to the Montal-Mueller method, where bilayers are formed between two experimental chambers (*cis* and *trans*) separated by a vertical 25 μm-thick Teflon layer with a 100 – 150 µm diameter aperture (**Figure 1**).^[7,8]^ The formation of a membrane is confirmed by measuring its capacitance, using two Ag/AgCl electrodes placed on each side of the membrane. For details about the protocol of membrane formation with the tested amphiphiles, see the Experimental section.

**Figure 1.**
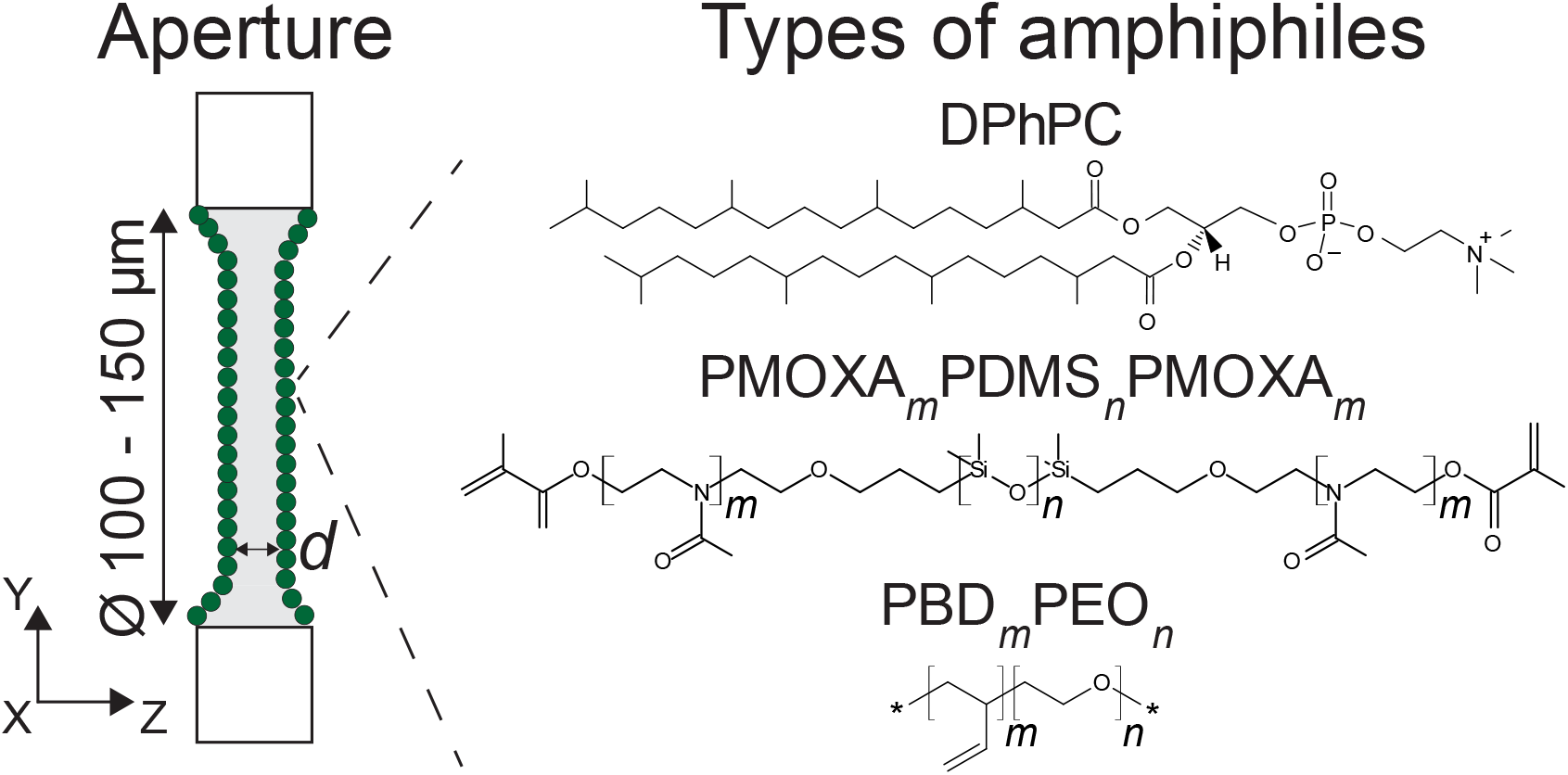
Components for free-standing membrane formation. On the left is a schematic drawing of the setup used in this work, showing a vertical hydrophobic substrate containing an aperture of 100 – 150 µm in diameter. Amphipathic membranes are then formed using the Montal-Mueller method.^[7]^ On the right are the amphiphiles used in this work: DPhPC (1,2-diphytanoyl-sn-glycero-3-phosphocoline), PMOXA_m_-PDMS_n_-PMOXA_m_ (poly(2-methyloxazoline-b-dimethylsiloxane-b-2-methyloxazoline), and PBD_m_-PEO_n_ (poly(butadiene)-b-ethylene oxide), where the parameters *m* and *n* are used to indicate the degree of polymerization. Depending on the amphiphile used, membranes can obtain a varying hydrophobic thickness *d*.

In our experimental chambers, DPhPC lipids (846 g/mol, with a hydrophobic bilayer thickness, *d*, 2.8 mn)^[29]^ yielded capacitance values of 150 – 170 pF. Membranes formed with PMOXA_4_PDMS_22_PMOXA_4_ (2280 g/mol) and PMOXA_18_PDMS_68_PMOXA_18_ (8000 g/mol) produced capacitance values of 100 – 120 pF and 70 – 80 pF, respectively (see Experimental section for details on choice of triblock polymers), reflecting the difference in membrane thicknesses. We estimated a hydrophobic thickness of 5 ≤ *d* ≤ 5.5 nm for PMOXA_4_PDMS_22_PMOXA_4_ (based on reported values for PMOXA_3_PDMS_19_PMOXA_3_ and PMOXA_4_PDMS_25_PMOXA_4_)^[30]^, while for PMOXA_18_PDMS_68_PMOXA_18_ *d* is reported to be 10 nm.^[14]^ In our systems, PMOXA_4_PDMS_22_PMOXA_4_ membranes were prone to rupture spontaneously under low applied potential, which may be enhanced by the inclusion of oil in our membrane formation protocol.^[31]^ However, excluding oil during the formation step did otherwise not allow membrane formation.

We also tested PBD_11_PEO_8_ (950 g/mol) diblock polymer bilayers, which were previously reported to successfully reconstitute α-HL and MspA nanopores.^[24–26]^ We estimated *d* ≈ 3.5 nm for PBD_11_PEO_8_ bilayers, assuming a slightly lower hydrophobic thickness than PBD_12_PEO_9_ (*d* = 3.7 nm).^[32]^ PBD_22_PEO_14_ polymers (1800 g/mol), forming thick bilayer membranes (*d* = 6.6 nm)^[33]^, were included as these membranes are known to still incorporate membrane proteins despite the large hydrophobic mismatch.^[31,34]^ In our setup, membranes formed with PBD-PEO polymers were easily formed and re-formed, yielding capacitance values of approximately 80 – 100 pF for PBD_22_PEO_14_ and 100 – 120 pF for PBD_11_PEO_8_. Again, the larger hydrophobic thickness of PBD_22_PEO_14_ compared to PBD_11_PEO_8_ is reflected by its lower capacitance value.

#### Electrical and biochemical stability of amphiphilic polymers

Under low applied potentials (i.e. +/- 100 mV), DPhPC membranes are stable for hours. As the voltage was gradually increased (for details of the protocol see the Experimental section), an average rupture potential of 240 ± 40 mV (average ± standard deviation) was measured (**Table 1**, N = 5), in agreement with previous studies.^[35,36]^ PBD_22_PEO_14_ showed the strongest voltage stability (rupture potential = 540 ± 50 mV), followed by PBD_11_PEO_8_ (rupture potential = 420 ± 60 mV). PMOXA_18_PDMS_68_PMOXA_18_ showed an average rupture potential similar to DPhPC (270 ± 30 mV). PMOXA_4_PDMS_22_PMOXA_4_ formed membranes that were prone to rupture spontaneously, even with low or no bias.

The biochemical stability of the polymer membranes was assessed by adding 25% v/v human serum directly to the experimental chambers (Table 1, see Experimental section and Figure S1 for details of the protocol). While DPhPC membranes consistently ruptured instantaneously under these conditions, both PBD_22_PEO_14_ and PBD_11_PEO_8_ (N = 10) remained stable upon the addition of serum while applying +150 mV for 15 minutes. PMOXA_18_PDMS_68_PMOXA_18_ also showed a higher biochemical stability than DPhPC, with 46% of the formed membranes (N = 13) withstanding the challenging conditions for the full 15 minutes under +150 mV (39% of the membranes ruptured within 5 minutes, 15% between 5 and 15 minutes). PMOXA_4_PDMS_22_PMOXA_4_ was not regarded due to its propensity to spontaneously rupture.

**Table 1.**
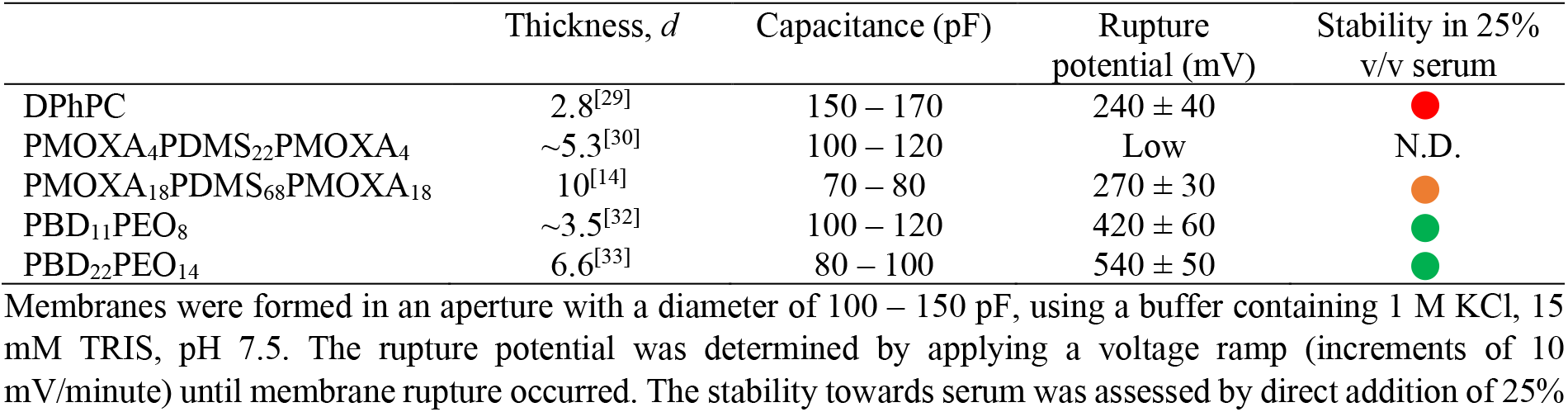
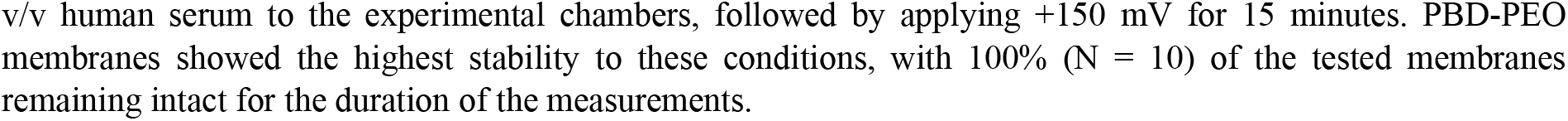
Properties of planar membranes formed from amphiphiles tested in this work.

| | Thickness, $d$ | Capacitance (pF) | Rupture potential (mV) | Stability in 25% v/v serum |
| --- | --- | --- | --- | --- |
| DPhPC | 2.8<sup>[29]</sup> | 150 – 170 | $240 \pm 40$ | ● |
| PMOXA<sub>4</sub>PDMS<sub>22</sub>PMOXA<sub>4</sub> | $\sim 5.3$<sup>[30]</sup> | 100 – 120 | Low | N.D. |
| PMOXA<sub>18</sub>PDMS<sub>68</sub>PMOXA<sub>18</sub> | 10<sup>[14]</sup> | 70 – 80 | $270 \pm 30$ | ● |
| PBD<sub>11</sub>PEO<sub>8</sub> | $\sim 3.5$<sup>[32]</sup> | 100 – 120 | $420 \pm 60$ | ● |
| PBD<sub>22</sub>PEO<sub>14</sub> | 6.6<sup>[33]</sup> | 80 – 100 | $540 \pm 50$ | ● |

Membranes were formed in an aperture with a diameter of 100 – 150 μm, using a buffer containing 1 M KCl, 15 mM TRIS, pH 7.5. The rupture potential was determined by applying a voltage ramp (increments of 10 mV/minute) until membrane rupture occurred. The stability towards serum was assessed by direct addition of 25%
v/v human serum to the experimental chambers, followed by applying +150 mV for 15 minutes. PBD-PEO membranes showed the highest stability to these conditions, with 100% (N = 10) of the tested membranes remaining intact for the duration of the measurements.

### Nanopore insertion and characterization in polymer membranes

Pore insertion in polymer membranes was tested by adding preformed oligomeric nanopores into solution under a constant applied potential, using buffer conditions typically used in nanopore experiments (1 M KCl, 15 mM TRIS, pH 7.5 for β-barrel nanopores α-HL, Cytotoxin K - CytK, MspA-M2, lysenin, and 150 mM NaCl, 15 mM TRIS, pH 7.5 for α-helical nanopores Cytolysin A - ClyA and YaxAB). Successful pore insertion was visualized by step-wise current increases. Pore reconstitution into thick polymer membranes could be achieved due to a voltage-induced membrane thinning effect, known as electrocompression, the magnitude of which scales quadratically with the applied voltage.^[37]^

The produced open pore current of a nanopore in a polymer membrane was averaged (<*I*_O_>) and compared to its corresponding value in DPhPC membranes to assess any changes in the pore’s ability to conduct ions (N = 3, see Experimental methods for details). β-barrel nanopores were measured at +75 mV, while the α-helical pores ClyA and YaxAB were measured at -75 mV, their typical working potential. The obtained current recordings also enabled to compare power spectra of nanopores in polymer membranes with similar spectra taken with DPhPC membranes, to analyze any shifts in electrical noise.

#### Nanopore insertion in PMOXA-PDMS-PMOXA membranes

Pore insertion was typically achievable within 15 minutes (while applying a constant potential in the ± 100 – 200 mV range) by adding up to 10× more pore oligomers into solution than with DPhPC. Upon insertion, all tested β-barrels nanopores (α-HL, lysenin, and MspA-M2) consistently obtained undesirable current properties not observed with DPhPC bilayers under the same conditions: substantial variability in the induced ionic currents; unstable current baselines; short current blockades attributed to pore gating; and (strongly) increased noise levels (Figure S2). All three nanopores showed large differences in <*I*_O_> values upon reconstitution in PMOXA-PDMS-PMOXA membranes relative to DPhPC bilayers (**Table 2**). Moreover, power spectra obtained from current signals could show large shifts in amplitude levels, indicating a strong increase in electrical noise (Figure S3 – S5). Finally, α-helical pores ClyA and YaxAB were also tested but were found to easily trigger membrane rupture upon insertion, preventing nanopore characterization.

**Table 2.** Percentage change of average open pore current values of nanopores in PMOXA-PDMS-PMOXA and PBD-PEO membranes compared to DPhPC membranes.

| | $\Delta\langle I_o \rangle$<br>PMOXA<sub>4</sub><br>PDMS<sub>22</sub><br>PMOXA<sub>4</sub> | $\Delta\langle I_o \rangle$<br>PMOXA<sub>18</sub><br>PDMS<sub>68</sub><br>PMOXA<sub>18</sub> | $\Delta\langle I_o \rangle$<br>PBD<sub>11</sub>PEO<sub>8</sub> | $\Delta\langle I_o \rangle$<br>PBD<sub>22</sub>PEO<sub>14</sub> | $\Delta\langle I_o \rangle$<br>PBD<sub>11</sub>PEO<sub>8</sub> +<br>DPhPC<br>(1:1 w/w) | $\Delta\langle I_o \rangle$<br>PBD<sub>22</sub>PEO<sub>14</sub> +<br>DPhPC<br>(2:1 w/w) |
| --- | --- | --- | --- | --- | --- | --- |
| α-HL<sup>a)</sup> | +32.8% ● | -29.8% ● | +0.7% ● | +15.2% ● | -6.1% ● | -9.8 % ● |
| Lysenin<sup>a)</sup> | -61.7% ● | -48.9% ● | -24.2% ● | N.D. | -6.0% ● | -11.9% ● |
| MspA-M2<sup>a)</sup> | -18.8% ● | -10.3% ● | -10.0% ● | N.D. | -4.7% ● | -3.1% ● |
| CytK<sup>a)</sup> | N.D. | N.D. | -21.8% ● | +14.2% ● | -5.7% ● | -14.5% ● |
| AeL<sup>a)</sup> | N.D. | N.D. | -22.6% ● | N.D. | +2.6% ● | +7.1% ● |
| ClyA<sup>b)</sup> | N.D. | N.D. | -58.8% ● | -61.7% ● | -11.8% ● | -29.7% ● |
| YaxAB<sup>b)</sup> | N.D. | N.D. | N.D. | N.D. | -15.8% ● | N.D. |

<sup>a)</sup> Experiments performed with: 75 mV, 1 M KCl, 15 mM TRIS, pH 7.5. <sup>b)</sup> Experiments performed with: -75 mV, 150 mM NaCl, 15 mM TRIS, pH 7.5. The traffic light colors next to $\Delta\langle I_O \rangle$ values are added to indicate the quality of the produced current signal of a nanopore in a corresponding membrane (i.e. observed noise level, baseline stability, level of pore gating). Green color indicates current properties similar to those in DPhPC membranes. Note: PBD<sub>11</sub>PEO<sub>8</sub> yielded good current signals with β-barrel nanopores, but nanopores were regularly observed to exit the membrane. Data was obtained from triplicates (N = 3) using a 10 kHz sample frequency and 2 kHz low-pass Bessel filter. See Table S2 for all averaged current values used for the current comparisons in this table.

#### Nanopore insertion in PBD-PEO membranes

In PBD_22_PEO_14_ (*d* = 6.6 nm), successful pore characterization was achieved for α-HL, cytotoxin K (CytK), and ClyA. However, reconstituted nanopores were observed to yield current signals with unstable baselines (Figure S2) and observable changes in <*I*_O_> values (Table 2). The insertion frequency for nanopores was very low: 10^3^× higher protein concentrations in solution relative to DPhPC and applied potentials up to +/-400 mV generally induced pore insertion once per several hours. The α-helical pore ClyA showed a strongly reduced open pore current, suggesting a highly constricted pore conformation. Current signals from α-HL and ClyA yielded power spectra with higher amplitude levels, indicating higher electrical noise (Figure S3, S6, and S7). Moreover, pore characterization in PBD_22_PEO_14_ was hindered by a common tendency of pores to exit the membrane shortly after reconstitution, a behavior rarely seen with DPhPC membranes. Ultimately, this led to the unsuccessful characterization of aerolysin (AeL) and MspA-M2, despite observed insertions (Figure S8). The attempted insertion of the α-helical pore YaxAB into PBD_22_PEO_14_ was unsuccessful, regardless of protein concentration.

Nanopore insertion in PBD_11_PEO_8_ (*d* ≈ 3.5 nm) bilayer membranes was already reported for α-HL and MspA-M2.^[24–26]^ Here, the β-barrel pores CytK, AeL, lysenin, and α-helical pore ClyA were also found to insert (Figure S2). Typically, pore insertion in PBD_11_PEO_8_ could be achieved within 15 minutes, by using up to 100× higher pore concentrations as for DPhPC and applying up to +/-250 mV. All pores were found to yield stable current baselines with no enhanced pore gating. However, PBD_11_PEO_8_ could have a substantial reducing effect on <*I*_O_> values (Table 2). This effect was again particularly observable for the α-helical pore ClyA, suggesting that PBD-PEO membranes in general may not be suitable for this nanopore. Power spectra typically yielded similar or slightly increased amplitude levels compared to DPhPC, except for ClyA, which increased by 3 orders in magnitude (Figure S3 – S7, S9). As with PBD_22_PEO_14_, the nanopores were still regularly observed to exit from PBD_11_PEO_8_ shortly after insertion (Figure S10). This suggests a lack of stabilizing nanopore-membrane interactions once nanopores are reconstituted in PBD-PEO membranes. This is a crucial shortcoming, severely limiting their use for nanopore recordings. Finally, insertion of YaxAB nanopores into PBD_11_PEO_8_ was attempted, but similar to PBD_22_PEO_14_ these attempts were unsuccessful, regardless of protein concentrations used.

In conclusion, we found that the triblock membrane PMOXA-PDMS-PMOXA is unsuitable for nanopore reconstitution, regardless of membrane thickness. The insertion of β-barrel nanopores α-HL, lysenin and MspA-M2 induced undesirable current properties (not observed in DPhPC membranes) while insertion of α-helical nanopores was found to induce membrane rupture. PBD-PEO bilayers showed improved current properties of reconstituted nanopores compared to PMOXA-PDMS-PMOXA membranes, especially PBD_11_PEO_8_ bilayers. Reconstituted β-barrel nanopores could show low-noise, stable current baselines with no enhanced gating behavior, although the characteristic open pore current could be reduced. However, a major drawback that was regularly observed with β-barrel nanopores was the exit of the nanopore from the membrane, showing a lack of stabilization of the nanopore by the membrane environment. Moreover, PBD_11_PEO_8_ was found to be unsuitable for the tested α-helical nanopores.

### Hybrid membrane stability

To improve the properties of the amphipathic membranes, we tested mixtures between PBD-PEO and DPhPC. Studies with hybrid polymersomes containing PC lipids in the fluid phase and PBD-PEO revealed well-mixed, stable membranes.^[33,38–40]^ Furthermore, an environment of 50% PBD-PEO polymers and 50% PC lipids has been shown to form an optimal hybrid membrane for the membrane protein cytochrome *bo*_3_, while also being suitable for multiple ABC transporters.^[34,41,42]^

For our study, hybrid mixtures of PBD_11_PEO_8_ + DPhPC (mixed at a 1:1 w/w) and PBD_22_PEO_14_ + DPhPC (mixed at 2:1 w/w) were investigated, resulting in membranes with roughly equal numbers of lipids and polymers. Hybrid membranes yielded rupture potentials (540 ± 110 mV and 420 ± 100 mV, for PBD_22_PEO_14_ and PBD_11_PEO_8_ hybrid membranes, respectively, N = 5) and capacitance values similar to pure PBD_11_PEO_8_ or PBD_22_PEO_14_ membranes, although the rapture potential showed a larger spread. Membranes could be easily reformed after induced membrane rupture. Similar to PBD-PEO membranes, 100% of the hybrid mixtures remained intact after the introduction of 25% v/v human serum and application of +150 mV for 15 minutes (see Table S1, N = 10 for both membranes).

### Nanopore insertion and characterization in hybrid membranes

#### Nanopore insertion in [PBD_11_PEO_8_ + DPhPC] membranes

Insertion of nanopores into [PBD_11_PEO_8_ + DPhPC] was obtained by applying +/-200 mV. Rewardingly, all tested nanopores were able to insert in stable pore conformations, producing stable, low-noise current signals (**Figure 2**). Protein concentrations used to obtain single pore insertions were generally 1 – 10× higher than for DPhPC membranes (except YaxAB requiring a hundredfold higher), enabling pore insertion typically within 15 minutes.

**Figure 2.**
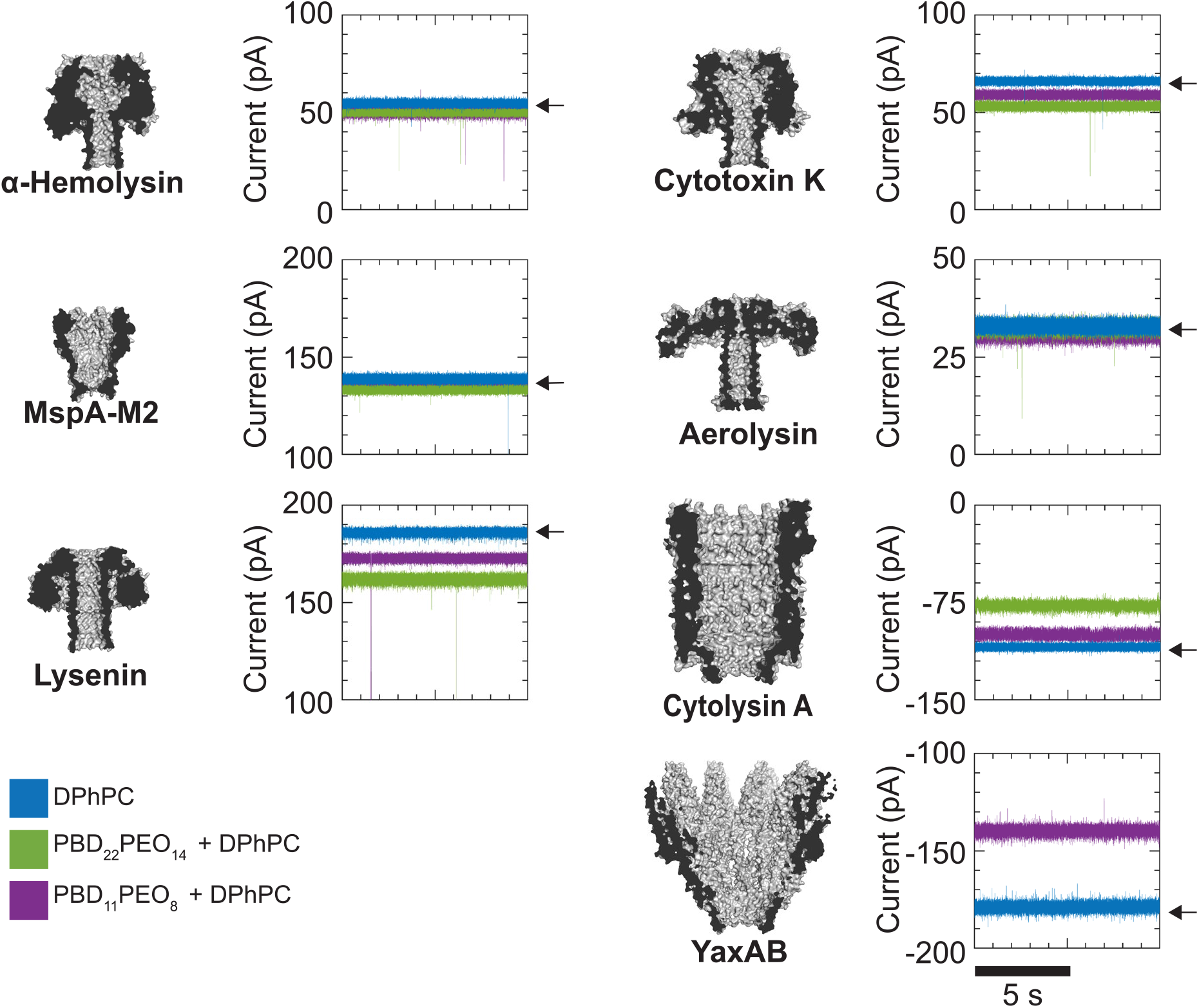
Electrical properties of nanopores in [PBD-PEO + DPhPC] membranes. Recordings in DPhPC, indicated by a small arrow, are included as a reference. For α-HL, MspA-M2, and AeL, traces for DPhPC, [PBD_11_PEO_8_ + DPhPC] and [PBD_22_PEO_14_ + DPhPC] are found to overlap. Conditions for all pores, except ClyA and YaxAB, included +75 mV, 1 M KCl, 15 mM TRIS, pH 7.5. ClyA and YaxAB were tested at conditions typically used in nanopore experiments: -75 mV, 150 mM NaCl, 15 mM TRIS, pH 7.5. Recordings were performed using a 10 kHz sampling frequency and a 2 kHz Bessel low-pass filter.

The effect of [PBD_11_PEO_8_ + DPhPC] on <*I*_O_> was relatively small for all pores (Table 2), with the largest changes observed for the α-helical pores ClyA (-11.8%) and YaxAB (-15.8%). Power spectra for pores in [PBD_11_PEO_8_ + DPhPC] were similar to those in DPhPC membranes, except for ClyA, showing a 10-fold increase in amplitude levels (Figure S3 – S7, S9, S11). An important observation with [PBD_11_PEO_8_ + DPhPC] was that pores did not show the tendency to exit the membrane, suggesting crucial stabilization by the presence of lipids.

#### Nanopore insertion in [PBD_22_PEO_14_ + DPhPC] membranes

Pore reconstitution in [PBD_22_PEO_14_ + DPhPC], mixed at 2:1 w/w, was stimulated with +/-250 mV. All pores, except YaxAB, could be successfully reconstituted (Figure 2). To maintain a pore insertion rate of approximately once per 15 minutes, MspA-M2 would require similar protein concentrations in solution as with DPhPC, while other nanopores needed a thousandfold higher concentration.

Compared to nanopores inserted into DPhPC membranes, [PBD_22_PEO_14_ + DPhPC] membranes had a small effect on <*I*_O_> for MspA-M2, while for other nanopores <*I*_O_> was more affected than with [PBD_11_PEO_8_ + DPhPC] (Table 2). Power spectra revealed that only ClyA showed a clear shift along the frequency spectrum towards higher magnitudes (Figure S3 – S7, S9). Finally, [PBD_22_PEO_14_ + DPhPC] also showed no early pore exit behavior, strengthening the hypothesis that lipids are vital for long-term membrane insertion.

### Molecular level organization of hybrid [PBD_11_PEO_8_ + DPhPC] membranes

To elucidate the origin of why hybrid membranes show the most preferable behavior, we investigated their molecular-level organization using molecular dynamics simulations. We chose to focus on the most successful hybrid membrane namely [PBD_11_PEO_8_ + DPhPC] at an equal lipid-to-polymer ratio. In particular, we used the coarse-grained (CG) Martini 3 force field, which previously has been applied to study nanopore systems.^[43–45]^ All simulations are started from a randomized initial mixture and are simulated for 6 µs after a short equilibration phase. We analyzed the homogeneity of the membranes by computing the normalized conditional entropy of mixing.^[46]^ This entropy is normalized such that a value of 100% corresponds to ideal mixing. By computing the normalized conditional entropy of mixing for a completely phase-separated system, which was built to yield the same system size, we can establish the lower bound to be about 20%. We note, that due to finite size effects in MD simulations, the lower bound is system size-dependent.

The normalized conditional entropy of mixing decreases as a function of simulation time for all three simulation replicas until it plateaus at a value of about 70% (**Figure 3**A). Thus, [PBD_11_PEO_8_ + DPhPC] hybrid membranes show a significant degree of phase separation, however, they do not reach a maximally phase-separated state. Previous molecular dynamics studies and experimental measurements have established that PBD_11_PEO_8_ forms homogeneously mixed membranes in the presence of lipids such as POPC and DOPC.^[47]^ Therefore, as a negative control, we also simulated mixtures with both these lipids at the Martini level resolution (Figure 3A). As expected, the resulting membranes are near to ideally mixed. Comparing the three lipids we see that the DPhPC-containing system shows the largest degree of non-ideal mixing. To further validate these results for [PBD_11_PEO_8_ + DPhPC] membranes and by extension the Martini model, we simulated three replicas of a smaller system using the all-atom CHARMM force field. These all-atom simulations show the same trend and a plateau entropy of mixing that is in good agreement with Martini simulations of the same size (Supporting Figure S12).

**Figure 3.**
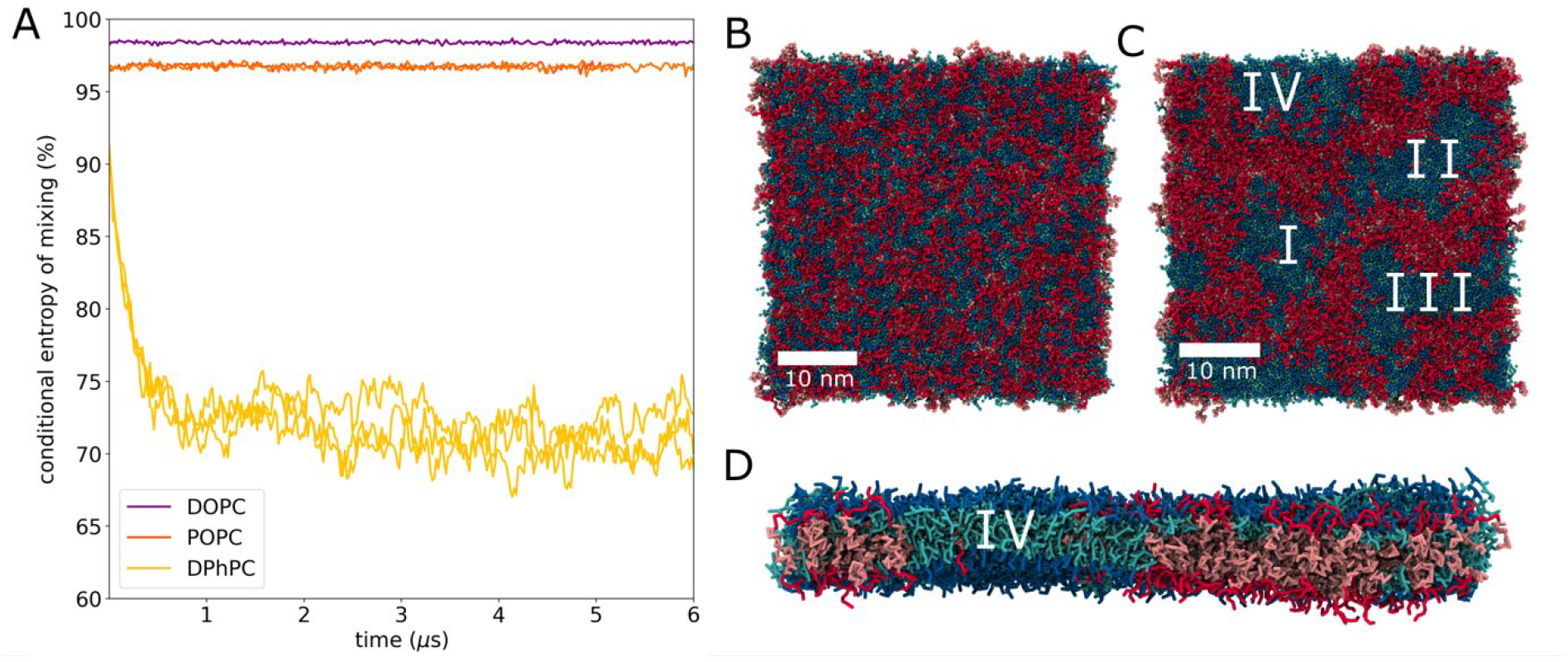
Mixing of [PBD_11_PEO_8_ + DPhPC] hybrid membranes. A) Conditional entropy of mixing computed for three replicas of [PBD_11_PEO_8_ + DPhPC] hybrid membranes (yellow) and two replicas each for [PBD_11_PEO_8_ + POPC] (orange) and [PBD_11_PEO_8_ + DOPC] hybrid membranes (purple). B) Initial frame of a 40 nm x 40 nm hybrid membrane with DPhPC. C) The same system as in B after 6 μs of simulation. Four lipid domains labeled I-IV are observed. D) Side view of the system in C. For systems B-D, lipid head groups are shown in blue and tails in cyan. PEO is shown in red, and the remaining polymer is shown in salmon pink.

**Figures 3B** and C show a snapshot of one of the DPhPC containing replicas at the beginning of the simulation and after 6 μs. One can see that the lipids form small domains (I-IV). These domains are dispersed throughout a segregated but mostly in-contact polymer matrix. Panel D shows the side view bisecting such a domain. To quantify the difference in phase separation between the lipids and polymers we performed a bin-based clustering of the molecules. Supplementary Figure S13 shows that the polymer molecules form a single cluster across all replicas, whereas the lipid clusters fluctuate in size. A similar picture is obtained when calculating the size of the largest lipid cluster (Supplementary Figure S14). In all replicas, the largest lipid cluster contains anywhere between 20%-70% of the total number of lipids molecules. Using this clustering method, we can approximately define the size of the four domains shown in Figure 3C. They contain on average 560 lipids and have an area of about 12 nm × 12 nm. The largest observed domains contain approximately 1500 lipids resulting in a domain size of about 16.7 nm. All analyses point towards the mixed membrane existing as an inhomogeneous system with lipid nanodomains interspersed in a polymer matrix.

To answer how the membrane organizes in the presence of the pores, we embedded three different nanopores (α-Hemolysin, Lysenin, and Aerolysin, using structural data from their corresponding PBD files 7ahl, 5gaq and 5jzt, respectively) in an initially mixed hybrid membrane and let the system self-organize. Each CG simulation was run in triplicate for 6 μs. For computational efficiency, we only simulated the barrels of the pores. **Figure 4A** shows an exemplary snapshot at the end of one of the replica simulations of the α-Hemolysin barrel. The top and side views reveal that the pore is surrounded by a lipid shell embedded in the lipid nanodomain.

**Figure 4.**
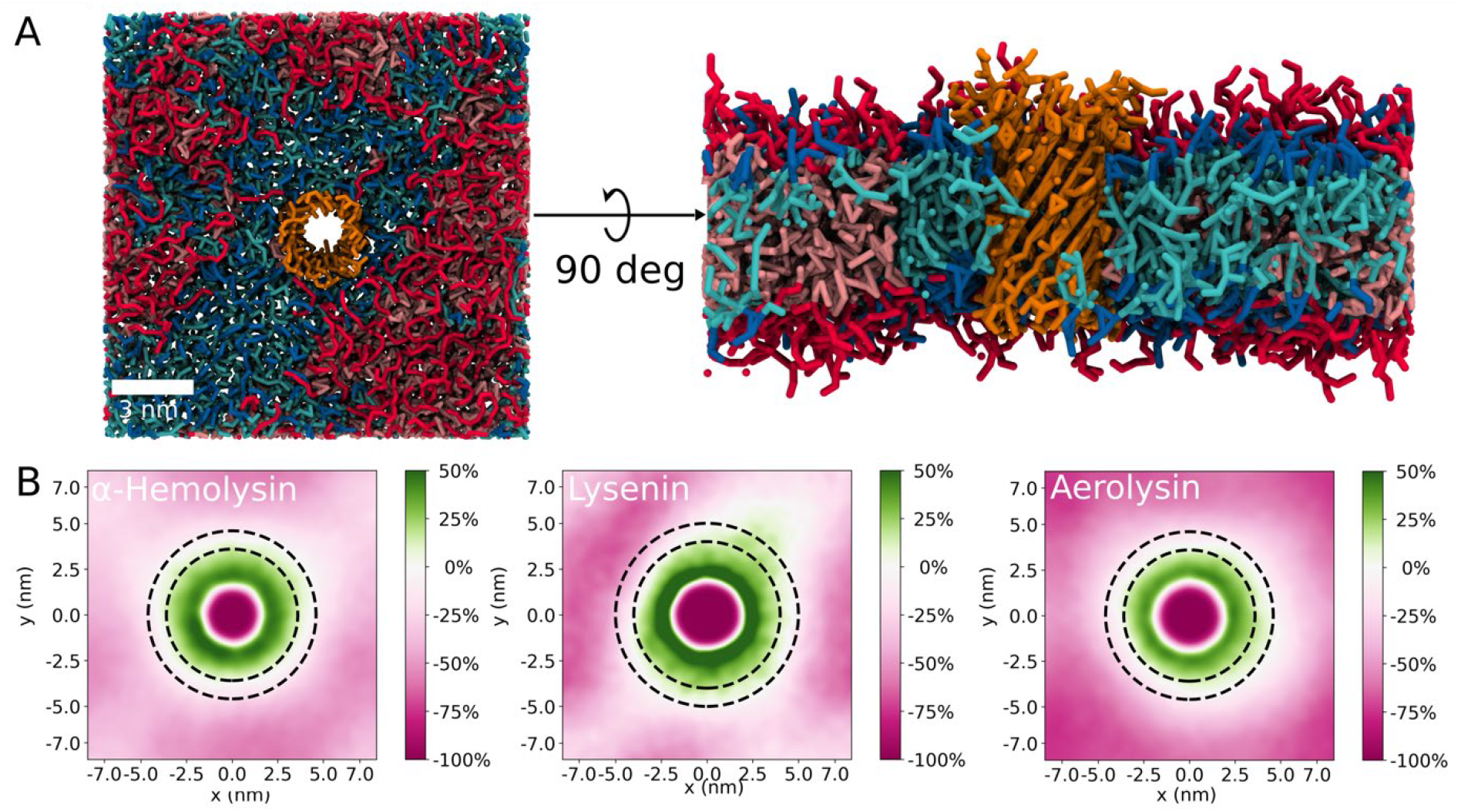
Nanopore proteins in hybrid membranes. A) Top view and side view of *α*-Hemolysin barrel embedded into a [PBD_11_PEO_8_ + DPhPC] hybrid membrane. B) Lipid enrichment around three nanopores. Only the upper leaflet is shown. Enrichments for all leaflets and replicas are shown in Supplementary Figure 15. Lipid enrichment is measured relative to the lipid density found within the area marked by the two dashed rings. Green areas correspond to a positive enrichment (i.e. more lipids) and the pink areas to a depletion of lipids (i.e. fewer lipids). As expected, there is 100% depletion inside the pores as there are no lipids present in the channels.

To further quantify membrane protein interactions, the lipid enrichment was computed. The enrichment describes how much more likely it is to find lipids at a distance from the protein relative to the density observed in a shell between 2.2 nm – 3.2 nm from the protein (indicated by dashed lines). **Figure 4B** shows the enrichment plots computed for one replica and centered around the protein. We define the upper leaflet to be where the head of the protein would be located. An enrichment is indicated in green whereas depletion is shown in pink. The pink area in the center of the plot corresponds to the channel, where lipids are 100% depleted as expected. In agreement with **Figure 4A** we can see a green halo surrounding the channel region, which means that the proteins are surrounded by lipids (i.e. located inside a nanodomain). Interestingly, we see a strong enrichment of up to 50% in the immediate vicinity across both leaflets and all pore types (Supplementary Figure 15).

### Single-molecule detection with biological nanopores in hybrid [PBD_11_PEO_8_ + DPhPC] membranes

To probe the biosensing ability of these pores, we carried out single-molecule measurements, using four different nanopores in [PBD_11_PEO_8_ + DPhPC] membranes, comparing results with similar measurements in DPhPC membranes (N = 3 for all measurements, Table 3).

γ-cyclodextrin (**Figure 5A**, γ-CD, 5 μM, *trans*, -100 mV) induced blockades of comparable scale to α-HL in the hybrid and lipid membranes, with increased dwell time (τ) but equal excluded current (*I*_ex_) values in the hybrid membrane (**Table 3**). Note that *I*_ex_ is defined here as: 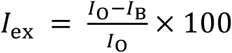, with *I*_B_ the blocked current value.

**Table 3.** Results of single-molecule detection in [PBD_11_PEO_8_ + DPhPC] membranes compared to DPhPC membranes.

|  | DPhPC | PBD<sub>11</sub>PEO<sub>8</sub> + DPhPC |
| --- | --- | --- |
| Analyte | Parameter tested | Parameter tested |
| γ-CD with α-HL<sup>a)</sup> | $\tau$: $537 \pm 46$ ms<br>$I_{ex}$: $52.3 \pm 1.1\%$<br>$n = 208 \pm 31$ | $\tau$: $656 \pm 85$ ms<br>$I_{ex}$: $53.2 \pm 1.1\%$<br>$n = 125 \pm 31$ |
| Biotinylated DNA 25-mer with MspA-M2<sup>b)</sup> | $\tau$: $0.88 \pm 0.07$ ms<br>$I_{ex}$: $46.4 \pm 0.4\%$<br>$n = 690 \pm 51$ | $\tau$: $0.89 \pm 0.07$ ms<br>$I_{ex}$: $46.4 \pm 0.3\%$<br>$n = 540 \pm 208$ |
| S1 peptide with CytK<sup>c)</sup> | $\tau$: $1.59 \pm 0.36$ ms<br>$I_{ex}$: $85.5 \pm 2.4\%$<br>$n = 547 \pm 259$ | $\tau$: $0.97 \pm 0.04$ ms<br>$I_{ex}$: $74.9 \pm 10.6\%$<br>$n = 543 \pm 196$ |
| SA and CRP with YaxAB<sup>d)</sup> | SA:<br>$\tau$: $188 \pm 15$ ms<br>$I_{ex}$: $50.96 \pm 2.4\%$<br>$n = 192 \pm 41$ | SA:<br>$\tau$: $14.76 \pm 2.98$ ms<br>$I_{ex}$: $51.1 \pm 0.5\%$<br>$n = 340 \pm 44.90$ |
| | CRP:<br>$\tau$: $52.3 \pm 7.9$ ms<br>$I_{ex}$: $21.87 \pm 0.58\%$<br>$n = 164 \pm 31$ | CRP:<br>$\tau$: $4.69 \pm 0.72$ ms<br>$I_{ex}$: $21.3 \pm 2.5\%$<br>$n = 272 \pm 48$ |

<sup>a)</sup> 5 μM γ-CD, *trans*, 100 mV, recording time 5 minutes. <sup>b)</sup> 0.5 μM, *cis*, +150 mV, recording time 100 sec. <sup>c)</sup> 300 nM S1 (123 amino acids, net charge +28 at pH 7.5), *cis*, -150 mV, recording time 90 sec. <sup>d)</sup> 0.25 μM SA + 0.25 μM CRP, *cis*, -75 mV, recording time 80 sec. γ-CD ( α-HL), biotinylated DNA (MspA-M2) and S1 (CytK) measurements were performed in 1 M KCl, 15 mM TRIS, pH 7.5. SA and CRP (YaxAB) were detected in a buffer containing 150 mM NaCl, 15 mM TRIS, pH 7.5. Measurements were performed in triplicate, yielding $n$ (average ± S.D.) events. Differences in $n$ values between measurements with DPhPC and [PBD<sub>11</sub>PEO<sub>8</sub> + DPhPC] membranes can be indicative to change in event frequency. Measurements were recorded with 50 kHz sample frequency and a 10 kHz Bessel low-pass filter.

**Figure 5.**
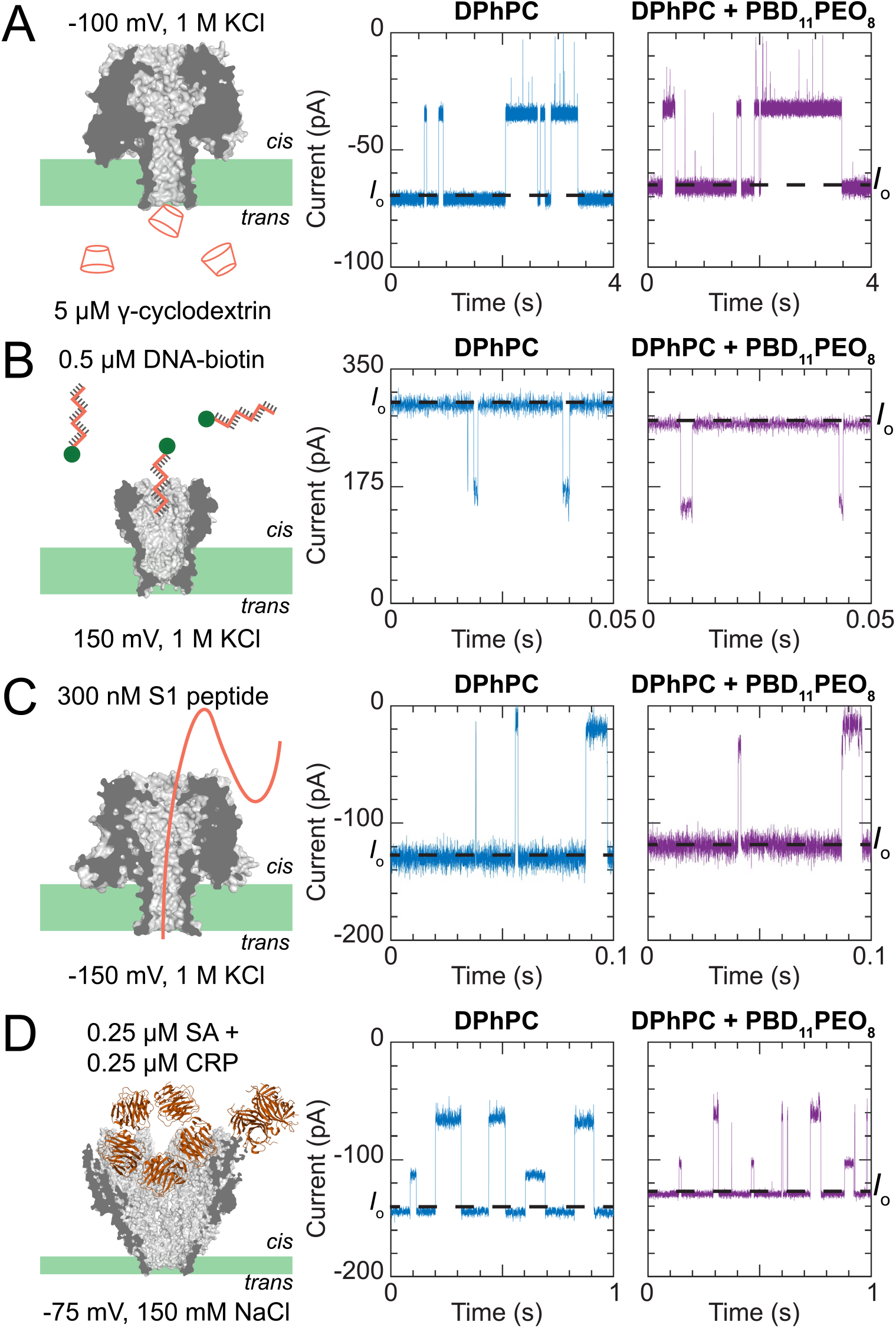
Single-molecule nanopore sensing in DPhPC and hybrid [PBD_11_PEO_8_ + DPhPC] membranes. A) Recordings of γ-cyclodextrin (γ-CD 5 µM, *trans*, -100 mV) binding to α-hemolysin. B) Translocation of 25-mer DNA-biotin (0.5 µM, *cis, +*150 mV) through MspA-M2. C) S1 peptide (300 nM, *cis, -*150 mV) translocation through CytK. D) Detection of streptavidin (SA, 53 kDa, 0.25 µM, *cis*) and C-reactive protein (CRP, 125 kDa, 0.25 µM, *cis*) by YaxAB at - 75 mV. The sampling frequency was set at 50 kHz with a 10 kHz lowpass Bessel filter. For presentation purposes, traces for B, C and D were also digitally filtered with a 2 kHz Gaussian filter. The buffer was 1 M KCl (panels A – C) or 150 mM NaCl (panel D), 15 mM TRIS, pH 7.5.

The translocation of a biotinylated 25-mer DNA (Figure 5B, 0.5 μM, *cis*, +150 mV) by MspA-M2 showed very similar current blockades for both τ and *I*_ex_ in [PBD_11_PEO_8_ + DPhPC] and DPhPC membranes (Table 3). As observed earlier with lipid membranes, the addition of streptavidin (SA, 25 nM, *cis*) also led to long DNA-biotin:SA blockade signals with MspA-M2 in [PBD_11_PEO_8_ + DPhPC] (Figure S16).^[48]^

CytK nanopores were tested for the translocation of model polypeptide ‘S1’, containing 123 amino acids and a net charge of +28 at pH 7.5, designed to remain unstructured in solution.^[49]^ When S1 (300 nM) was added to the *cis* chamber (Figure 5C, -150 mV), blockades with longer dwell time and lower *I*_ex_ values were observed with CytK in [PBD_11_PEO_8_ + DPhPC] membranes compared to the same events measured in DPhPC membranes (Table 3). The slight differences might reflect small changes in the pore conformation, affecting pore-substrate interactions.

Finally, the effect of the membrane on the α-helical YaxAB was tested. A mixture of SA (53 kDa) and C-reactive protein (CRP, 125 kDa), which were previously used, were added to the *cis*-side (-75 mV, 0.25 µM of each protein).^[5]^ The resulting blockade levels for SA and CRP showed similar *I*_ex_ values in [DPhPC + PBD_11_PEO_8_] membranes as in DPhPC membranes, whereas τ was reduced in the hybrid membrane (Table 3). However, the resulting dwell time vs. *I*_ex_ scatter plot still reveals two distinguishable clusters for SA and CRP in [PBD_11_PEO_8_ + DPhPC] membranes, similar to what is observed with DPhPC membranes (see Figure S17). This is an indication that the conical conformation of YaxAB is retained in the hybrid membrane, allowing the identification of differently sized proteins in hybrid membranes.

### Stable single-molecule detection of human serum with biological nanopores in hybrid [PBD_11_PEO_8_ + DPhPC] membranes

The addition of 1% v/v human serum to the aqueous environment of reconstituted nanopores (ClyA: *cis*, -40 mV; YaxAB: *cis*, -50 mV; lysenin: *trans*, +100 mV) enabled single-molecule recordings (**Figure 6A – C**) without observing short-term membrane rupture (Figure S18). By contrast, similar measurements in DPhPC could only be performed for a few minutes.^[5]^ Higher serum concentrations could not be tested for ClyA and YaxAB because of the high frequency of blockades. Owing to its narrower diameter, lysenin showed fewer and shorter current blockades when serum dilutions were added to the *trans* side. When serum was added to the *cis*-side, no current blockades were observed (neither at +100 mV or - 100 mV). Hence, to illustrate that measurements in high concentrations of serum are possible, we tested a lysenin nanopore exposed to a solution with 25% v/v human serum added to both *cis* and *trans* compartments. Under this high concentration of serum, many long-lasting blockades were observed, reflecting the entry of multiple serum components into the pore (Figure 6C). Most importantly, it remained possible to perform stable recordings without experiencing instantaneous membrane rupture as observed with DPhPC bilayers (10 min. stable recording with no sign of membrane degradation shown in Figure S19).

**Figure 6.**
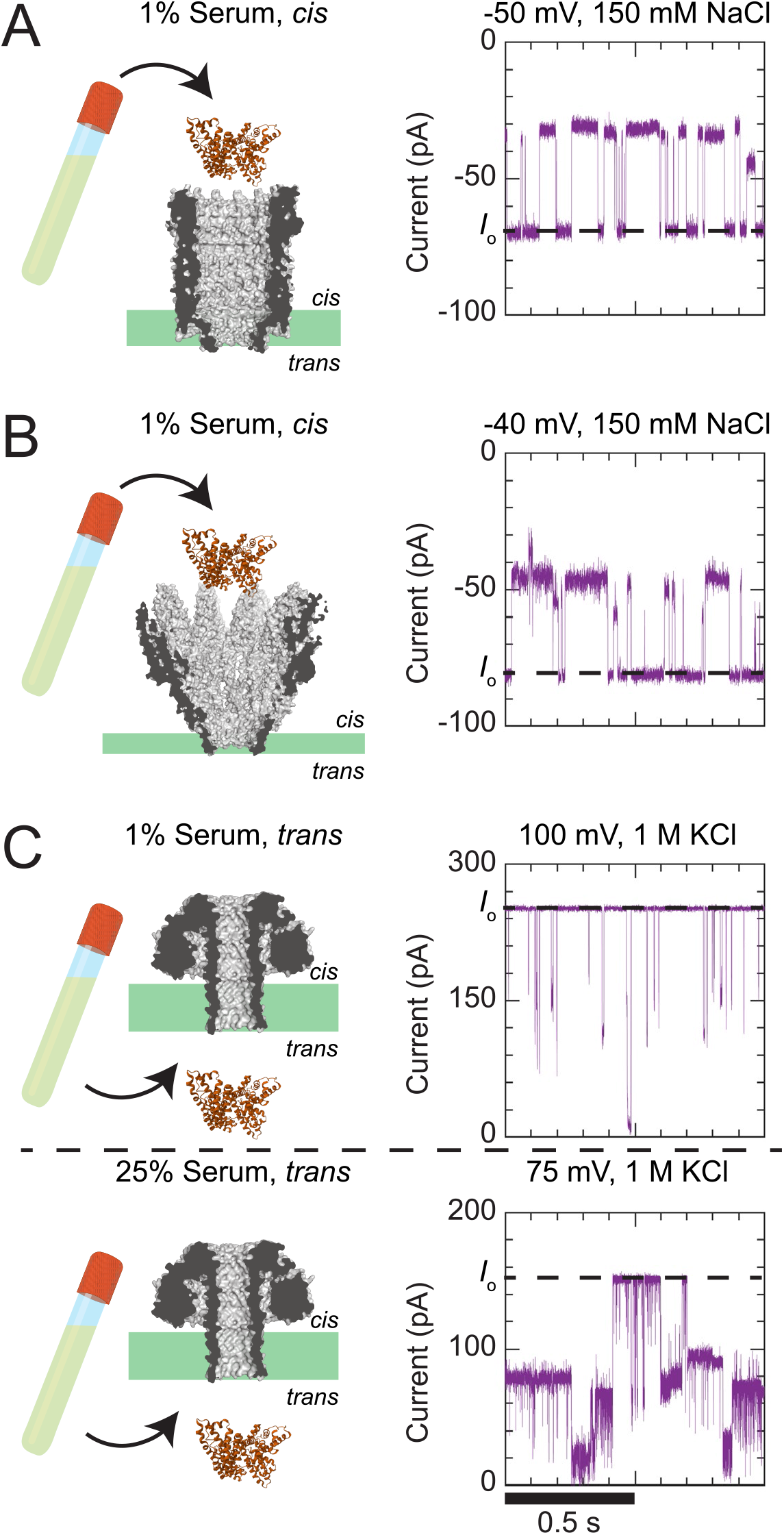
Single-molecule recordings of human serum with ClyA, YaxAB, and lysenin in [PBD_11_PEO_8_ + DPhPC] membranes. A) Events from 1% serum added to the *cis* side with a ClyA nanopore at -50 mV (150 mM NaCl, 15 mM TRIS, and pH 7.5). B) Events from 1% serum (in 150 mM NaCl, 15 mM TRIS, pH 7.5) with YaxAB at -40 mV. C) The top trace shows events from 1% serum (*trans*, +100 mV, 1 M KCl, 15 mM TRIS, pH 7.5) detected with lysenin. If serum was added to *cis*, no events were observed at either +100 mV or -100 mV. The bottom current trace shows events from 25% serum (*cis* + *trans*, +75 mV) diluted in a buffer containing 1 M KCl, 15 mM TRIS, pH 7.5.

Overall, these results indicate that biological nanopores in hybrid [PBD_11_PEO_8_ + DPhPC] membranes can form a stable interface for single-molecule detection of human serum, paving the way for novel nanopore-based biosensors capable of measuring complex biological samples.

## Conclusions

In recent years, nanopores have emerged as a reliable tool for the detection of molecules from biological samples, showcasing their potential as single-molecule sensors. However, in laboratory settings nanopores are typically inserted in lipid bilayer membranes. These membranes are fragile and prone to rupture when interfaced with biological samples, hindering the development of nanopore-based biosensors in commercial devices, or when using complex biological samples.^[5,6]^

Here, we have shown that membranes consisting of amphiphilic polymers can offer improved stability towards human serum. Specifically, diblock PBD-PEO polymers possessed superior stability properties in the presence of high concentrations of human serum and applied electric potentials. However, pure PBD-PEO bilayers were found to be unsuitable for the tested α-helical pores, such as ClyA (highly constricted upon reconstitution) and YaxAB no reconstitution observed). Further, the reconstitution of β-barrel nanopores could present several undesirable observations: strongly reduced insertion rates; early membrane exits; unstable baselines; increased electrical noise or large changes in open pore current values.

Nonetheless, we found that hybrid membranes formed with approximately a 50:50 molar distribution of PBD-PEO polymers and DPhPC lipids can retain the favorable membrane stability of pure PBD-PEO membranes while allowing electrical recordings comparable to pure DPhPC membranes. Hybrid membranes of PBD_11_PEO_8_ and DPhPC (mixed at 1:1 w/w) are found to form the most suitable environment for the various tested nanopores. The hybrid membrane environment allowed for successful insertion of all tested nanopores, regardless of pore size and/or shape, or whether the nanopores contained α-helical or β-barrel transmembrane domains. Importantly, the presence of lipids is likely to be crucial to preventing nanopores from exiting the membrane prematurely. Molecular simulations revealed that at a 1:1 molar composition, the amphipathic molecules mixed non-homogeneously with lipids forming nanodomains interspersed by a polymer matrix. Nanopores were then observed to partition within the lipid nanodomains and sequester lipids to their immediate surroundings. Hence, these hybrid membranes allow mimicking the natural lipid environment around the nanopore, as demonstrated by the insertion of α-helical and β-barrel nanopore alike, and the comparable single-molecule nanopore measurements recorded in DPhPC and hybrid membranes. At the same time, the hybrid polymer-lipid membranes enabled stable detection of proteins from human serum well above the limit of lipid membranes, indicating that such hybrid membranes largely retain their stability advantages of the pure polymer membranes.

The results presented in this study resolve a serious challenge for the analysis of biological fluids with nanopores, opening up new routes for novel biosensors.

## Experimental section

### Chemicals

The following block polymers were purchased from Polymer Source, Inc.: PMOXA_4_PDMS_22_PMOXA_4_ (Poly(2-methyloxazoline-b-dimethylsiloxane-b-2-methyloxazoline, Mn (g/mol): 330-b-1620-b-330, P42401F-MEOXZDMSMEOXZ); PMOXA_18_PDMS_68_PMOXA_18_ (Methacrylate End Functionalized Poly(2-methyloxazoline-b-dimethylsiloxane-b-2-methyloxazoline, Mn (g/mol): 1.5-b-5.0-b-1.5, P18536A-MAMOXZDMSMOXZMA); PBD_22_PEO_14_ ((Poly)butadiene-b-ethylene oxide, Mn (g/mol): 1200-b-600, P41745-BdEO); PBD_11_PEO_8_ ((Poly)butadiene-b-ethylene oxide, Mn (g/mol): 600-b-350, P41807C-BdEO). Note: PMOXA_18_PDMS_68_PMOXA_18_ contained methacrylate moieties at both polymer ends that can allow UV-crosslinking, however, this was not exploited. DPhPC (1,2-diphytanoyl-sn-glycero-3-phosphocoline, CAS#207131-40-6), hexadecane 99% (CAS#544-76-3), chloroform 99% (CAS#67-66-3), γ-cyclodextrin (γ-CD, CAS#7585-39-9), C-reactive protein (CRP, AG723) and human serum (H4522-100ML) were ordered from Sigma-Aldrich. 1 µM γ-CD stock suspension was prepared with MilliQ, samples were stored at 4 °C. CRP was purchased in lyophilized state, MilliQ was added to obtain a 25 µM stock solution that was stored at 4 °C. Pentane 99% (CAS%109-66-0) and Streptavidin (SA, 21122) were ordered from ThermoFisher Scientific. SA stock solutions were prepared by resuspension in MilliQ to 50 µM (determined via Bradford assay) and samples were stored at 4 °C. Sodium chloride (CAS#7647-14-5), Potassium chloride (CAS#7447-40-7) and TRIS (CAS#77-86-1) were acquired from Roth. DNA-biotin 25-mer was purchased from Integrated DNA Technologies (IDT) with the following sequence: /5’-biotin/AAAAATAATACGACTGACTATAGGG. DNA-biotin was acquired in lyophilized state, MilliQ was used to prepare 100 µM stock suspensions and storage was at 4 °C for short-term use. S1 peptide was prepared in our laboratory by Adina Sauciuc and the protocol is thoroughly described in literature.^[53]^ The concentration of the prepared stock sample was determined as 1.73 mg/mL via Bradford assay.

### Expression, purification, and oligomerization of nanopores

Detailed protocols used for the preparation of nanopores are well described in previous work: CytK, AeL, ClyA-AS, and YaxAB.^[5,49–51]^ For α-HL, the protocol for CytK was followed. For Lysenin, the protocol developed for FraC was followed.^[52]^

In short, the pT7sc1 plasmid containing the AeL, α-HL, ClyA, CytK, Lysenin, and MspA-M2 constructs, and the pRSET-A plasmid containing the YaxA and YaxB constructs were transformed into *E. cloni* ®EXPRESS BL21(DE3) electrocompetent cells. All constructs were cultured in 400 mL 2×YT medium supplemented with 100 μg/mL ampicillin at 37 °C and 200 rpm. Protein expression was induced when the OD_600_ of the culture reached 0.8, by addition of 0.5 mM Isopropyl β-D-thiogalactopyranoside (IPTG) and subsequent overnight culturing at 25 °C and 200 rpm.

The cell pellets of AeL, ClyA, and Lysenin were resuspended in lysis buffer containing 50 mM Tris-HCl pH 7.5, 150 mM NaCl, and 20 mM imidazole.^[50,51]^ The cells were disrupted via probe sonication (Brandson) and the protein was purified with Ni-NTA affinity chromatography (Qiagen). The nanopores were eluted from the beads with 3× 250 µL fractions of elution buffer containing 50 mM Tris-HCl pH 7.5, 150 mM NaCl, and 300 mM imidazole.

The cell pellets containing CytK and α-HL were resuspended in lysis buffer containing 50 mM HEPES, pH 7.5, 150 mM NaCl, 20 mM imidazole, 0.02% DDM, and one tablet of protease inhibitor EDTA-free per cell pellet of 100 mL culture.^[49]^ The proteins were purified by Ni-NTA affinity chromatography and eluted from the beads with 3× 250 µL fractions of elution buffer containing 50 mM Tris-HCl, pH 7.5, 150 mM NaCl, 250 mM imidazole, and 0.02% DDM.

The cell pellets of MspA-M2 were resuspended in lysis buffer containing 50 mM Tris-HCl pH 7.5, 150 mM NaCl, 0.02% DDM. The protein was purified by *Strep*Tactin affinity chromatography and eluted from the beads with 3× 250 µL fractions of elution buffer containing 50 mM Tris-HCl, pH 8.5, 150 mM NaCl, 5 mM *d*-Desthiobiotin and 0.02% DDM. The cell pellets containing YaxA and YaxB were resuspended in lysis buffer containing 50 mM Tris-HCl pH 8.0, 300 mM NaCl, 20 mM imidazole (2 M urea for YaxA), and one tablet of protease inhibitor EDTA-free.^[5]^ The protein was purified by Ni-NTA affinity chromatography and eluted from the beads with 3× 250 µL fractions of elution buffer containing 50 mM Tris-HCl, pH 8.0, 300 mM NaCl and 100 mM imidazole.

Oligomerization of α-HL, CytK, and MspA-M2 occurred during purification with 0.02% DDM. For oligomerization of the other nanopores, 1 mg/mL protein concentrations were used. Oligomerization of AeL was triggered by cleaving the pro-peptide with trypsin for 15 minutes (1:1000 trypsin:protein ratio), which creates the active form of AeL. ClyA was oligomerized by addition of 0.2% DDM and 30 minutes incubation at 37 °C. Different oligomeric forms of ClyA were separated on a Native-PAGE gel (4 – 20%, Criterion Bio-rad). Lysenin monomers were oligomerized on DPhPC:sphingomyelin liposomes (1:1 ratio, 10 mg/mL), by incubation for 30 minutes at 37 °C in a 1:10 protein:liposome ratio. The liposomes were then dissolved by incubating 5 minutes with 0.6% N,N-dimethyldodecylamine-N-oxide (LDAO). The oligomers were diluted in 20 mL buffer (50 mM Tris-HCl, pH 7.5, 150 mM NaCl and 0.02% DDM). The lysenin oligomers were further purified with a second Ni-NTA affinity purification and eluted from the beads with 200 µL of elution buffer containing 50 mM Tris-HCl pH 7.5, 150 mM NaCl, 200 mM EDTA and 0.02% DDM. YaxAB oligomerization was triggered by incubating YaxA and YaxB together in a 1:1 ratio for 30 minutes. The acquired YaxAB oligodimers were separated by incubating with 1.5% 6-cyclohexylhexyl β-d-maltoside (Cymal-6) for 30 minutes at 4 °C. The YaxAB oligomers were further purified with size exclusion chromatography on the Superose6 10/300 GL Increase (Cytiva) column pre-equilibrated with SEC buffer containing 25 mM HEPES pH 7.0, 150 mM NaCl, 0.05% Cymal-6.

### Membrane formation

Membranes were formed in an in-house fabricated device. The device consisted of two separate chambers (700 µL capacity each), mounted together. The chambers had adjacent orifices, that were separated by a 25-µm thick Teflon sheet.^[8]^ Membranes were formed in an aperture (diameter 100 – 150 µm) located in the Teflon surface. The protocol for membrane formation with DPhPC and PBD-PEO slightly differed from that for PMOXA-PDMS-PMOXA. DPhPC and PBD-PEO were dissolved in pentane (5 and 10 mg/mL, respectively). With these amphiphiles, the Teflon substrate was first wetted with 5 µL of 4% v/v hexadecane in pentane. 400 µL of a buffer solution was then added to each chamber. Finally, 20 µL of the amphiphile solution was added to the buffer-air interface and the pentane was allowed to evaporate. PMOXA-PDMS-PMOXA polymers were dissolved in chloroform due to its poor solubility in pentane (10 mg/mL). Since chloroform is less volatile than pentane, the polymer solution was first added to the dry chambers to ensure complete evaporation of the solvent (20 µL per chamber for PMOXA_4_PDMS_22_PMOXA_4_, 50 µL per chamber for PMOXA_18_PDMS_68_PMOXA_18_ to ensure an excess of polymers). After the chloroform had evaporated, the Teflon layer was wetted with 5 µL of 4% v/v hexadecane in pentane. 400 µL of buffer was then added to each chamber. The formation of the membrane was induced by subsequent lowering and raising of the buffer level past the aperture, using a pipette. The successful formation of a membrane was deduced from an increase in membrane capacitance. Ag/AgCl electrodes were inserted into each chamber, the *trans* chamber contained the working electrode, and the *cis* chamber the grounding electrode. The formation of a bilayer could be confirmed by voltage-induced rupture (1.3 V), followed by the formation of a new membrane. This control step was easily achievable for DPhPC, PBD-PEO, and PMOXA_18_PDMS_68_PMOXA_18_ membranes. PMOXA_4_PDMS_22_PMOXA_4_ membranes typically could not be reformed after rupture due to its overall mechanical instability.

### Selection of triblock polymers

When selecting PMOXA-PDMS-PMOXA polymers (**Figure 1**), care was taken to select polymers with a hydrophilic weight fraction (*f*_hydrophilic_) within 25 – 45% to prevent the formation of micelles or other unwanted aggregates: PMOXA_4_PDMS_22_PMOXA_4_ (2280 g/mol, *f*_hydrophilic_ = 28.9%) and PMOXA_18_PDMS_68_PMOXA_18_ (8000 g/mol, *f*_hydrophilic_ = 37.5%).^[30]^

### Protocols for assessing electrical and biochemical stability properties of amphilic membranes

The electrical stability properties of the amphiphilic membranes were characterized by their rupture potentials (N = 5). For each test, membranes were formed in a buffer containing 1 M KCl and 15 mM TRIS at pH = 7.5. Then, a voltage ramp (10 mV steps, each applied for 1 minute) was applied and the potential at which the membrane ruptured was recorded. Membrane stability towards biological fluids was assessed using human serum at a concentration greatly exceeding the limit for lipid membranes. First, membranes were formed with 300 μL of buffer in each chamber (1 M KCl, 15 mM TRIS, pH 7.5). Then, 100 μL of human serum was added to each chamber to obtain a final serum concentration of 25% v/v, and a constant potential of +150 mV was applied to the membrane for 15 minutes (Figure S1). The constant applied potential was briefly paused every 10 seconds to reconfirm the presence of a bilayer.

### Data acquisition and analysis

The electrophysiology setup used an Axopatch 200B patch clamp amplifier connected to a DigiData 1440 A/D converter. Experiments were monitored and analyzed with Clampex 10.7 (Molecular Devices) software. The current comparison measurements were performed with a sampling frequency of 10 kHz and a Bessel lowpass filter of 2 kHz. The averaged open pore current <*I*_O_> was taken from 3 individual measurements. For each measurement, an average *I*_O_ value was obtained from a Gaussian fit on a histogram of all pores’ current values taken in a 10-second timeframe (bin size 0.5 pA). These values were then averaged over the 3 measurements to obtain <*I*_O_>. The single-molecule measurements were carried out with a 50 kHz sampling frequency and 10 kHz Bessel lowpass filter. The current traces presented in Figure 5B and 5C were additionally digitally filtered with a Gaussian 2 kHz lowpass filter for presentation purposes. For excluded current (*I*_ex_) calculations, the open pore current (*I*_O_) was determined with a Gaussian fit on a histogram containing all current values (bin size 0.5 pA). The excluding current (*I*_ex_) was determined via: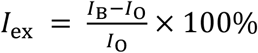, where *I*_B_ stands for the blocked current value obtained when an analyte is (partially) blocking the current during translocation. For event detection, events were included that induced a current blockade exceeding a threshold value. The threshold value corresponded to 5 times the standard error of the open pore current *I*_O_. For measuring γ-CD dwell times, since dwell time values could vary several orders of magnitude, events were grouped in histograms with logarithmic binning and fitted with a log probability exponential. For the detection of DNA-biotin and S1 peptide a conventional histogram was used (given the smaller spread of dwell times) and fitted with a standard exponential. Dwell time values with events from CRP and SA measurements were determined using a cumulative histogram and a standard exponential fitting, similar to our earlier work.^[5]^ The bin size of the histogram was chosen as at least three-fold smaller than the average dwell time obtained from the fitting. Measurements were performed in triplicate. The data obtained with γ-CD detection was taken from 5 minutes of measurement time and included an average of 208 ± 17 (DPhPC) and 125 ± 31 ([PBD_11_PEO_8_ + DPhPC]) data points. For DNA-biotin detection, data was acquired from 100 s measurement time and contained 735 ± 13 (DPhPC) and 540 ± 208 ([PBD_11_PEO_8_ + DPhPC]) data points. For S1 detection a time window of 90 seconds was taken, in which 547 ± 259 data points were taken with DPhPC and 543 ± 196 with [PBD_11_PEO_8_ + DPhPC] membranes. Finally, for analysis of SA and CRP measurements, a time interval of 80 s was used, resulting in 164 ± 32 events with DPhPC bilayers and 340 ± 45 events with [PBD_11_PEO_8_ + DPhPC] membranes.

### Molecular dynamics simulations

All molecular dynamics simulations were performed using GROMACS (version 2023 or 2023.4).^[53]^ Simulations at the all-atom level utilized the CHARMM36m force field (version July2022) for proteins and ions, as well as CHARMM TIP3P water.^[54,55]^ CHARMM parameters for DPhPC were published in the literature.^[56]^ PEO-b-PBD was modeled using the CHARMM general force field.^[57]^ After obtaining parameters for a small oligomer, the polyply python suite was used to create library files that allow the generating of arbitrarily long block copolymers.^[58]^ PEO-b-PBD parameters are available online (https://github.com/marrink-lab/polyply_1.0). Protein simulations of the nanopore proteins in pure DPhPC membranes were set up using the CHARMM-GUI membrane builder.^[59–62]^ Mixed hybrid membrane systems were created by back mapping from a Martini CG simulation using polyply.^[58]^ After equilibration, each simulation was run under constant temperature at 298.15 K using the v-rescale temperature coupling (τ = 1 ps) with a coupling group for solvent and polymer.^[63]^ The pressure was kept constant at 1 bar using the Parrinello–Rahman semi-isotropic pressure coupling algorithm (τ = 5 ps, β = 4.5 × 10^−5^ bar^−1^).^[64]^ Simulation lengths, system sizes, and compositions are listed in Supplementary Table S2.

Simulations at the CG level of resolution were performed using the Martini 3 force field.^[65]^ For all simulations, the SN3r-Q5 interaction level was changed to 3kJ/mol to improve PEO membrane association as previously suggested.^[66]^ Simulation parameters for hybrid membranes including parameters for DPhPC are shared in the zenodo repository. Protein parameters for α-Hemolysin and Lysenin were obtained using martinize2 starting from a reference protein structure obtained from a 300ns equilibration in DPhPC at the all-atom level.^[67]^ Protein parameters for Aerolysin were also generated using martinize2 but using a frame from a previously simulated system (ID: MD_aero_SMD).^[68]^ We note that residue GLU237 is considered protonated as was done in the reference simulation. Initial structures for membrane systems that did not contain proteins were generated using TS2CG, whereas those including proteins were created using a polyply building protocol.^[69]^ Simulation lengths, system sizes, and system compositions are listed in Supplementary Table S4. Run parameters were the same as recommended for Martini on GROMACS with special settings for large membranes (i.e. verlet-buffer-tolerance=-1 and rlist=1.35nm).^[65,70]^ After equilibration, each simulation was run under constant temperature at 298.15 K using the v-rescale temperature coupling (τ = 1 ps) with a coupling group for solvent, lipid, polymer, and protein if applicable.^[63]^ The pressure was kept constant at 1 bar using the Parrinello–Rahman semi-isotropic pressure coupling algorithm (τ = 12 ps, β = 3.0 × 10^−4^ bar^−1^).^[64]^

Membrane mixing was characterized by computing the conditional entropy of mixing following the ideas presented by Brandani and coworkers.^[46]^ However, instead of the nearest neighbor approximation the system was binned using a previously described procedure.^[71]^ Using this binned system, the entropy is obtained by considering the nearest neighbors in each direction except for diagonal neighbors. To determine the cluster sizes and their distribution, a similar binning approach was performed. Afterward, all bins that were in contact and fully surrounded by bins of the same type were clustered. Bins that have a different species as a neighbor are not considered to be part of a cluster but the periphery. This strategy is similar to a DBSCANclustering where interface points are discarded but using a binned system.^[72]^

Lipid densities around the proteins were computed by centering the trajectory around the protein and wrapping all atoms into the periodic box. Subsequently, the NC3 bead positions were replaced by a Gaussian of width 2.6 Å, and then the density was computed using the Freud library.^[73]^ Lipid protein contacts were determined as follows: A lipid is in contact with a specific protein residue if the NC3 bead, which represents the Choline head group in the Martini force field, is within a cut-off distance of 5.3 Å. The cut-off distance corresponds to the VdW minimum of a regular Martini bead. The number of contacts per residue is then averaged over the trajectory and subsequently over the chains and three replicas.

## Supporting Information

Supporting Information is available from the Wiley Online Library or from the author.

## Acknowledgments

We thank other members of the Giovanni Maglia lab for the generous sharing of nanopore oligomers and analytes, and C. Presutti for useful discussions and sharing of polymer samples. The experimental part of the project was financially sponsored by NWO VICI (no. 192068). The computational part is supported by funding from the ERC with the Advanced grant 101053661 “COMP-O-CELL”. F. G. acknowledges funding from the Klaus Tschira Foundation.

## Data availability statement

All experimental and computational data can be found at the corresponding Zenodo repository.

## Author Contributions

E.V., K.T. and G.M. designed the experiments, E.V. performed the experiments. N.J.H. expressed, purified and oligomerized all biological nanopores. E.V. and G.M. executed the experimental data analysis. F. G. designed, performed, and analyzed the molecular dynamics simulations. All authors contributed to the writing of the manuscript.

## References

[1] A. K. Thakur, L. Movileanu, ACS Sens 2019, 4, 2320.

[2] M. Ahmad, J.-H. Ha, L. A. Mayse, M. F. Presti, A. J. Wolfe, K. J. Moody, S. N. Loh, L. Movileanu, Nat Commun 2023, 14, 1374.

[3] N. S. Galenkamp, M. Soskine, J. Hermans, C. Wloka, G. Maglia, Nat Commun 2018, 9, 4085.

[4] G. Huang, A. Voorspoels, R. C. A. Versloot, N. J. van der Heide, E. Carlon, K. Willems, G. Maglia, Angew Chem Int Ed Engl 2022, 61, e202206227.

[5] S. Straathof, G. Di Muccio, M. Yelleswarapu, M. Alzate Banguero, C. Wloka, N. J. van der Heide, M. Chinappi, G. Maglia, ACS Nano 2023, 17, 13685.

[6] S. J. Greive, L. Bacri, B. Cressiot, J. Pelta, ACS Nano 2024, 18, 539.

[7] M. Montal, P. Mueller, Proc Natl Acad Sci U S A 1972, 69, 3561.

[8] G. Maglia, A. J. Heron, D. Stoddart, D. Japrung, H. Bayley, Methods Enzymol 2010, 475, 591.

[9] F. Bonté, R. L. Juliano, Chem Phys Lipids 1986, 40, 359.

[10] T. M. Allen, L. G. Cleland, Biochim Biophys Acta 1980, 597, 418.

[11] R. De, M. K. Mahata, K.-T. Kim, Adv Sci (Weinh) 2022, 9, e2105373.

[12] H. Cabral, K. Miyata, K. Osada, K. Kataoka, Chem Rev 2018, 118, 6844.

[13] C. Nardin, T. Hirt, J. Leukel, W. Meier, Langmuir 2000, 16, 1035.

[14] C. Nardin, M. Winterhalter, W. Meier, Langmuir 2000, 16, 7708.

[15] H. Bermudez, A. K. Brannan, D. A. Hammer, F. S. Bates, D. E. Discher, Macromolecules 2002, 35, 8203.

[16] M. P. Goertz, L. E. Marks, G. A. Montaño, ACS Nano 2012, 6, 1532.

[17] T. Einfalt, D. Witzigmann, C. Edlinger, S. Sieber, R. Goers, A. Najer, M. Spulber, O. Onaca-Fischer, J. Huwyler, C. G. Palivan, Nat Commun 2018, 9, 1127.

[18] A. Moquin, J. Ji, K. Neibert, F. M. Winnik, D. Maysinger, ACS Omega 2018, 3, 13882.

[19] P. J. Photos, L. Bacakova, B. Discher, F. S. Bates, D. E. Discher, J Control Release 2003, 90, 323.

[20] P. P. Ghoroghchian, P. R. Frail, K. Susumu, D. Blessington, A. K. Brannan, F. S. Bates, B. Chance, D. A. Hammer, M. J. Therien, Proc Natl Acad Sci U S A 2005, 102, 2922.

[21] W. Meier, C. Nardin, M. Winterhalter, Angew Chem Int Ed Engl 2000, 39, 4599.

[22] D. Wong, T.-J. Jeon, J. Schmidt, Nanotechnology 2006, 17, 3710.

[23] D. Morton, S. Mortezaei, S. Yemenicioglu, M. J. Isaacman, I. C. Nova, J. H. Gundlach, L. Theogarajan, J Mater Chem B 2015, 3, 5080.

[24] L. Yu, X. Kang, M. A. Alibakhshi, M. Pavlenok, M. Niederweis, M. Wanunu, Biophys J 2021, 120, 1537.

[25] M. Pavlenok, L. Yu, D. Herrmann, M. Wanunu, M. Niederweis, Biophys J 2022, 121, 742.

[26] X. Kang, C. Wu, M. A. Alibakhshi, X. Liu, L. Yu, D. R. Walt, M. Wanunu, ACS Nano 2023, 17, 5412.

[27] J. Habel, M. Hansen, S. Kynde, N. Larsen, S. R. Midtgaard, G. V. Jensen, J. Bomholt, A. Ogbonna, K. Almdal, A. Schulz, C. Hélix-Nielsen, Membranes (Basel) 2015, 5, 307.

[28] M. Garni, S. Thamboo, C.-A. Schoenenberger, C. G. Palivan, Biochim Biophys Acta Biomembr 2017, 1859, 619.

[29] N. Kučerka, M.-P. Nieh, J. Katsaras, Biochim Biophys Acta 2011, 1808, 2761.

[30] A. Najer, A. Belessiotis-Richards, H. Kim, C. Saunders, F. Fenaroli, C. Adrianus, J. Che, R. L. Tonkin, H. Høgset, S. Lörcher, M. Penna, S. G. Higgins, W. Meier, I. Yarovsky, M. M. Stevens, Small 2022, 18, e2201993.

[31] M. Kumar, J. E. O. Habel, Y. Shen, W. P. Meier, T. Walz, J Am Chem Soc 2012, 134, 18631.

[32] D. R. Barden, H. Vashisth, Faraday Discuss 2018, 209, 161.

[33] R. Seneviratne, G. Coates, Z. Xu, C. E. Cornell, R. F. Thompson, A. Sadeghpour, D. P. Maskell, L. J. C. Jeuken, M. Rappolt, P. A. Beales, Small 2023, 19, e2206267.

[34] S. Khan, M. Li, S. P. Muench, L. J. C. Jeuken, P. A. Beales, Chem Commun (Camb) 2016, 52, 11020.

[35] I. van Uitert, S. Le Gac, A. van den Berg, Biochim Biophys Acta 2010, 1798, 21.

[36] K. W. Langford, B. Penkov, I. M. Derrington, J. H. Gundlach, J Lipid Res 2011, 52, 272.

[37] N. Tamaddoni, G. Taylor, T. Hepburn, S. Michael Kilbey, S. A. Sarles, Soft Matter 2016, 12, 5096.

[38] J. Nam, P. A. Beales, T. K. Vanderlick, Langmuir 2011, 27, 1.

[39] R. Seneviratne, R. Catania, M. Rappolt, L. J. C. Jeuken, P. A. Beales, Soft Matter 2022, 18, 1294.

[40] W. A. Müller, P. A. Beales, A. R. Muniz, L. J. C. Jeuken, Biomacromolecules 2023, 24, 4156.

[41] R. Seneviratne, S. Khan, E. Moscrop, M. Rappolt, S. P. Muench, L. J. C. Jeuken, P. A. Beales, Methods 2018, 147, 142.

[42] S. Rottet, S. Iqbal, P. A. Beales, A. Lin, J. Lee, M. Rug, C. Scott, R. Callaghan, Polymers (Basel) 2020, 12, 1049.

[43] S. Zhang, G. Huang, R. C. A. Versloot, B. M. H. Bruininks, P. C. T. de Souza, S.-J. Marrink, G. Maglia, Nat Chem 2021, 13, 1192.

[44] V. Maingi, J. R. Burns, J. J. Uusitalo, S. Howorka, S. J. Marrink, M. S. P. Sansom, Nat Commun 2017, 8, 14784.

[45] M. Xue, L. Cheng, I. Faustino, W. Guo, S. J. Marrink, Biophys J 2018, 115, 494.

[46] G. B. Brandani, M. Schor, C. E. MacPhee, H. Grubmüller, U. Zachariae, D. Marenduzzo, PLoS ONE 2013, 8, e65617.

[47] Y. K. Go, C. Leal, Chem. Rev. 2021, 121, 13996.

[48] R. F. Purnell, K. K. Mehta, J. J. Schmidt, Nano Lett 2008, 8, 3029.

[49] A. Sauciuc, B. Morozzo Della Rocca, M. J. Tadema, M. Chinappi, G. Maglia, Nat Biotechnol 2023, DOI 10.1038/s41587-023-01954-x.

[50] R. C. A. Versloot, S. A. P. Straathof, G. Stouwie, M. J. Tadema, G. Maglia, ACS Nano 2022, 16, 7258.

[51] N. S. Galenkamp, V. Van Meervelt, N. L. Mutter, N. J. van der Heide, C. Wloka, G. Maglia, Methods Mol Biol 2021, 2186, 11.

[52] N. L. Mutter, G. Huang, N. J. van der Heide, F. L. R. Lucas, N. S. Galenkamp, G. Maglia, C. Wloka, Methods Mol Biol 2021, 2186, 3.

[53] M. J. Abraham, T. Murtola, R. Schulz, S. Páll, J. C. Smith, B. Hess, E. Lindahl, SoftwareX 2015, 1–2, 19.

[54] J. Huang, S. Rauscher, G. Nawrocki, T. Ran, M. Feig, B. L. De Groot, H. Grubmüller, A. D. MacKerell, Nat Methods 2017, 14, 71.

[55] W. L. Jorgensen, J. Chandrasekhar, J. D. Madura, R. W. Impey, M. L. Klein, The Journal of Chemical Physics 1983, 79, 926.

[56] X. Zhuang, J. R. Makover, W. Im, J. B. Klauda, Biochimica et Biophysica Acta (BBA) - Biomembranes 2014, 1838, 2520.

[57] K. Vanommeslaeghe, E. Hatcher, C. Acharya, S. Kundu, S. Zhong, J. Shim, E. Darian, O. Guvench, P. Lopes, I. Vorobyov, A. D. Mackerell, J Comput Chem 2010, 31, 671.

[58] F. Grünewald, R. Alessandri, P. C. Kroon, L. Monticelli, P. C. T. Souza, S. J. Marrink, Nat Commun 2022, 13, 68.

[59] S. Jo, T. Kim, W. Im, PLoS ONE 2007, 2, e880.

[60] E. L. Wu, X. Cheng, S. Jo, H. Rui, K. C. Song, E. M. Dávila-Contreras, Y. Qi, J. Lee, V. Monje-Galvan, R. M. Venable, J. B. Klauda, W. Im, J. Comput. Chem. 2014, 35, 1997.

[61] J. Lee, X. Cheng, J. M. Swails, M. S. Yeom, P. K. Eastman, J. A. Lemkul, S. Wei, J. Buckner, J. C. Jeong, Y. Qi, S. Jo, V. S. Pande, D. A. Case, C. L. Brooks, A. D. MacKerell, J. B. Klauda, W. Im, J. Chem. Theory Comput. 2016, 12, 405.

[62] S. Jo, T. Kim, V. G. Iyer, W. Im, J Comput Chem 2008, 29, 1859.

[63] G. Bussi, D. Donadio, M. Parrinello, The Journal of Chemical Physics 2007, 126, 014101.

[64] M. Parrinello, A. Rahman, Journal of Applied Physics 1981, 52, 7182.

[65] P. C. T. Souza, R. Alessandri, J. Barnoud, S. Thallmair, I. Faustino, F. Grünewald, I. Patmanidis, H. Abdizadeh, B. M. H. Bruininks, T. A. Wassenaar, P. C. Kroon, J. Melcr, V. Nieto, V. Corradi, H. M. Khan, J. Domański, M. Javanainen, H. Martinez-Seara, N. Reuter, R. B. Best, I. Vattulainen, L. Monticelli, X. Periole, D. P. Tieleman, A. H. De Vries, S. J. Marrink, Nat Methods 2021, 18, 382.

[66] F. Grünewald, Material Design Using Martini: Accelerating Discovery through Coarse-Grained Simulations, University of Groningen, 2023.

[67] P. C. Kroon, F. Grunewald, J. Barnoud, M. Van Tilburg, P. C. T. Souza, T. A. Wassenaar, S. J. Marrink, 2023, DOI 10.7554/eLife.90627.1.

[68] H. Ouldali, K. Sarthak, T. Ensslen, F. Piguet, P. Manivet, J. Pelta, J. C. Behrends, A. Aksimentiev, A. Oukhaled, Nat Biotechnol 2020, 38, 176.

[69] W. Pezeshkian, M. König, T. A. Wassenaar, S. J. Marrink, Nat Commun 2020, 11, 2296.

[70] H. Kim, B. Fábián, G. Hummer, J. Chem. Theory Comput. 2023, 19, 8919.

[71] M. P. A. Van Tilburg, S. J. Marrink, M. König, F. Grünewald, J. Chem. Theory Comput. 2024, 20, 212.

[72] M. Ester, H. P. Kriegel, J. Sander, X. Xiaowei, “A Density-Based Algorithm for Discovering Clusters in Large Spatial Databases with Noise,” can be found under https://www.aaai.org, 1996.

[73] V. Ramasubramani, B. D. Dice, E. S. Harper, M. P. Spellings, J. A. Anderson, S. C. Glotzer, Computer Physics Communications 2020, 254, 107275.

