## Supplementary information for "Hybrid lipid-block copolymer membranes enable stable reconstitution of a wide range of nanopores and robust sampling of serum"

F. Grünewald

Heidelberg Institute for Theoretical Studies (HITS), Heidelberg, Germany

### Tables

Table S1: **Membrane stability in the presence of Serum**. The membrane stability was tested by introducing serum on both chambers (25% final v/v), to the aqueous phase at each side of the membrane containing 1 M KCl, 15 mM TRIS, pH 7.5. A potential of +150 mV was then applied for 15 minutes to assess the chemical stability of each membrane to serum. DPhPC membrane ruptured within seconds, $PMOXA_4PDMS_{22}PMOXA_4$, were too unstable in buffer to perform measurements.

| **Membrane** | **Percentage of membranes that successfully completed 25% v/v serum stability test** |
|---|---|
| DPhPC | 0% - Instantaneous rupture |
| $PMOXA_4PDMS_{22}PMOXA_4$ | **-** |
| $PMOXA_{18}PDMS_{68}PMOXA_{18}$ | 46% (N =13) |
| $PBD_{11}PEO_8$ | 100% (N = 10) |
| $PBD_{22}PEO_{14}$ | 100% (N =10) |
| $PBD_{11}PEO_8$ + DPhPC 1:1 w/w | 100% (N =10) |
| $PBD_{22}PEO_{14}$ + DPhPC 2:1 w/w | 100% (N =10) |

Table S2 **Open pore current values of nanopores characterized in DPhPC, PMOXA-PDMS-PMOXA, and PBD-PEO membranes.**

| | | DPhPC | $PMOXA_mPDMS_nPMOXA_m$ | | $PBD_mPEO_n$ | |
|---|---|---|---|---|---|---|
| | | | $m = 4, n = 22$ | $m = 18, n = 68$ | $m = 11, n = 8$ | $m = 22, n = 14$ |
| α-HL[a)] | $\langle I_O \rangle \pm \sigma$ [pA] | 55.3 ± 0.5 | 89.0 ± 6.8 | 38.8 ± 15.7 | 55.7 ± 1.1 | 77.3 ± 7.3 |
| Lysenin[a)] | $\langle I_O \rangle \pm \sigma$ [pA] | 183.7 ± 1.6 | 70.4 ± 20.8 | 93.8 ± 12.9 | 139.3 ± 3.4 | N.D. |
| MspA-M2[a)] | $\langle I_O \rangle \pm \sigma$ [pA] | 137.6 ± 1.0 | 111.7 ± 24.5 | 123.4 ± 33.1 | 123.9 ± 2.9 | N.D. |
| CytK[a)] | $\langle I_O \rangle \pm \sigma$ [pA] | 62.9 ± 2.2 | N.D. | N.D. | 49.2 ± 1.6 | 71.8 ± 1.5 |
| AeL[a)] | $\langle I_O \rangle \pm \sigma$ [pA] | 30.8 ± 1.5 | N.D. | N.D. | 25.0 ± 0.3 | N.D. |
| ClyA[b)] | $\langle I_O \rangle \pm \sigma$ [pA] | -111.0 ± 4.3 | N.D. | N.D. | -45.6 ± 22.7 | -40.3 ± 7.5 |
| YaxAB[b)] | $\langle I_O \rangle \pm \sigma$ [pA] | -168.1 ± 7.8 | N.D. | N.D. | N.D. | N.D. |

[a)] Experiments performed with: 75 mV, 1 M KCl, 15 mM TRIS, pH 7.5. [b)] Experiments performed with: -75 mV, 150 mM NaCl, 15 mM TRIS, pH 7.5. Data was obtained from triplicates (N = 3) using a 10 kHz sample frequency and 2 kHz low-pass Bessel filter.

Table S3 **Open pore current properties of nanopores characterized in DPhPC and hybrid [PBD-PEO + DPhPC] membranes** [a]

| | | DPhPC | $PBD_{11}PEO_8$ + DPhPC (1:1 w/w) | $PBD_{22}PEO_{14}$ + DPhPC (2:1 w/w) |
|---|---|---|---|---|
| α-HL[a)] | $<I_O> \pm \sigma$ [pA] | 55.3 ± 0.5 | 51.7 ± 2.7 | 49.9 ± 0.3 |
| Lysenin[a)] | $<I_O> \pm \sigma$ [pA] | 183.7 ± 1.6 | 172.7 ± 2.3 | 161.9 ± 0.9 |
| MspA-M2[a)] | $<I_O> \pm \sigma$ [pA] | 137.6 ± 1.0 | 131.1 ± 5.1 | 133.3 ± 1.3 |
| CytK[a)] | $<I_O> \pm \sigma$ [pA] | 62.9 ± 2.2 | 59.3 ± 1.5 | 53.8 ± 0.9 |
| AeL[a)] | $<I_O> \pm \sigma$ [pA] | 30.8 ± 1.5 | 31.6 ± 1.1 | 33.0 ± 0.5 |
| ClyA[b)] | $<I_O> \pm \sigma$ [pA] | -111.0 ± 4.3 | -97.9 ± 2.5 | -78.0 ± 0.1 |
| YaxAB[b)] | $<I_O> \pm \sigma$ [pA] | -168.1 ± 7.8 | -141.6 ± 7.0 | N.D. |

[a)] Experiments performed with: 75 mV, 1 M KCl, 15 mM TRIS, pH 7.5. [b)] Experiments performed with: -75 mV, 150 mM NaCl, 15 mM TRIS, pH 7.5. Data was obtained from triplicates (N = 3) using a 10 kHz sample frequency and 2 kHz low-pass Bessel filter.

Table S4. **Specifications of systems simulated.** Tag refers to the folder in the Zenodo repository where the trajectories can be found.

| Tag | Force Field | LIP - Type | #LIP | POL - Type | #POL | Size (nm) | Protein | Length |
|---|---|---|---|---|---|---|---|---|
| RA1 | CHARMM36 | DPhPC | 475 | N/A | N/A | 14.5 x 14.5 | 5gaq | 300ns |
| RA2 | CHARMM36 | DPhPC | 421 | N/A | N/A | 14 x 14 | 7ahl | 300ns |
| MA1 | CHARMM36 | DPhPC | 134 | 8_11 | 134 | 10 x 10 | N/A | 6µs |
| MA2 | CHARMM36 | DPhPC | 134 | 8_11 | 134 | 10 x 10 | N/A | 6µs |
| MA3 | CHARMM36 | DPhPC | 134 | 8_11 | 134 | 10 x 10 | N/A | 6µs |
| MA4 | CHARMM36 | DPhPC | 134 | 8_11 | 134 | 10 x 10 | N/A | 7µs |
| MC1 | Martini3 | DPhPC | 134 | 8_11 | 134 | 10 x 10 | N/A | 6µs |
| MC6 | Martini3 | DPhPC | 2162 | 8_11 | 2162 | 40 x 40 | N/A | 6µs |
| MC7 | Martini3 | DPhPC | 2162 | 8_11 | 2162 | 40 x 40 | N/A | 7µs |
| MC8 | Martini3 | DPhPC | 2162 | 8_11 | 2162 | 40 x 40 | N/A | 7µs |
| MC9 | Martini3 | POPC | 2162 | 8_11 | 2162 | 40 x 40 | N/A | 6 µs |
| MC10 | Martini3 | POPC | 2162 | 8_11 | 2162 | 40 x 40 | N/A | 6 µs |
| MC11 | Martini3 | DOPC | 2162 | 8_11 | 2162 | 40 x 40 | N/A | 6 µs |
| MC12 | Martini3 | DOPC | 2162 | 8_11 | 2162 | 40 x 40 | N/A | 6 µs |
| PC1 | Martini3 | DPhPC | 334 | 8_11 | 334 | 17x17 | 7ahl | 6µs |
| PC2 | Martini3 | DPhPC | 334 | 8_11 | 334 | 17x17 | 7ahl | 6µs |
| PC3 | Martini3 | DPhPC | 334 | 8_11 | 334 | 17x17 | 7ahl | 6µs |
| PC4 | Martini3 | DPhPC | 334 | 8_11 | 334 | 17x17 | 5gaq | 6µs |
| PC5 | Martini3 | DPhPC | 334 | 8_11 | 334 | 17x17 | 5gaq | 6µs |
| PC6 | Martini3 | DPhPC | 334 | 8_11 | 334 | 17x17 | 5gaq | 6µs |
| PC7 | Martini3 | DPhPC | 334 | 8_11 | 334 | 17x17 | 5jzt | 6µs |
| PC8 | Martini3 | DPhPC | 334 | 8_11 | 334 | 17x17 | 5jzt | 6µs |
| PC9 | Martini3 | DPhPC | 334 | 8_11 | 334 | 17x17 | 5jzt | 6µs |

### Figures

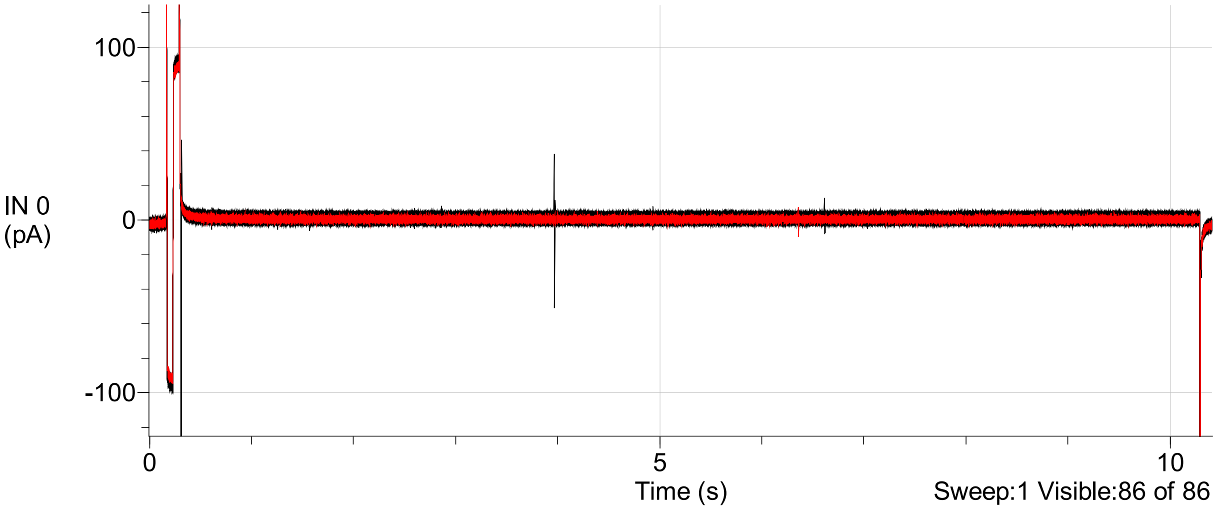

Figure S1: **Overview of voltage protocol applied during serum stability tests on membranes.** The protocol consisted of sweeps of approximately 11 seconds. During each sweep, first a brief triangular waveform was applied to confirm the presence of a membrane, followed by a constant applied voltage of +150 mV. For each individual measurement, the protocol was applied for 15 minutes.

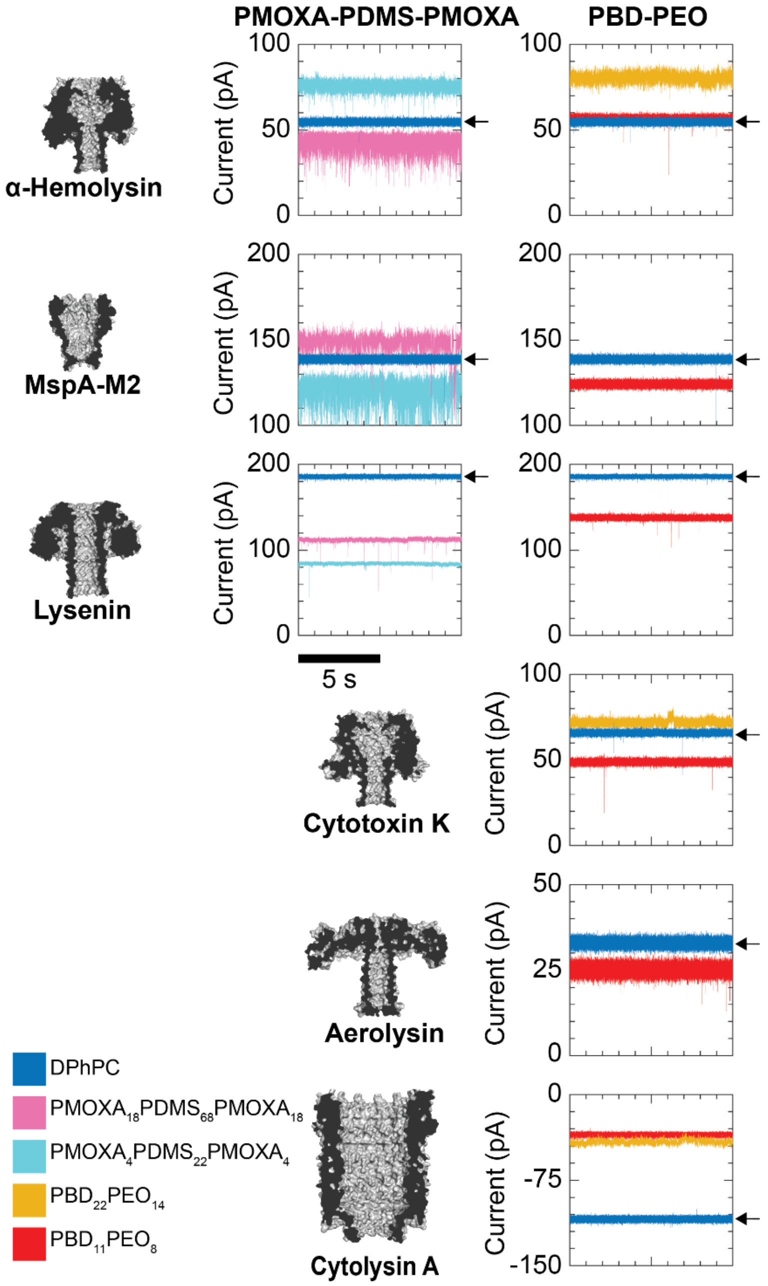

Figure S2: **Electrical properties of nanopores in polymer-based membranes.** Recordings in DPhPC, indicated by a small arrow, are included as a reference. Left column: recordings in PMOXA-PDMS-PMOXA membranes. Right column: recordings in PBD-PEO membranes. Although β-barrel nanopores inserted in $PBD_{11}PEO_8$ presented current properties comparable to those in DPhPC membranes, regular ejection from the membrane was observed (like in $PBD_{22}PEO_{14}$), creating a serious drawback of using $PBD_{11}PEO_8$ membranes for nanopore recordings. Conditions for all pores, except ClyA and YaxAB, included +75 mV, 1 M KCl, 15 mM TRIS, pH 7.5. ClyA and YaxAB were tested at conditions typically used in nanopore experiments: -75 mV, 150 mM NaCl, 15 mM TRIS, pH 7.5. Recordings were performed using a 10 kHz sampling frequency and a 2 kHz Bessel low-pass filter.

DPhPC

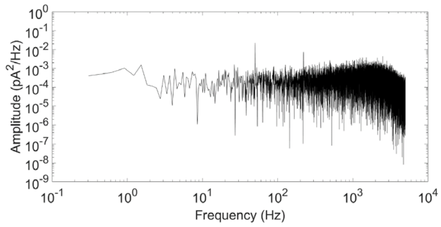

PMOXA4PDMS22PMOXA4

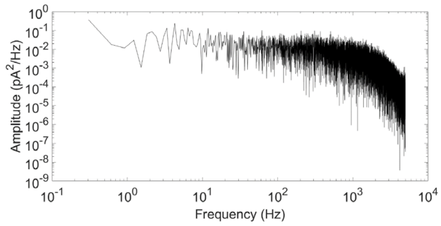

PMOXA18PDMS68PMOXA18

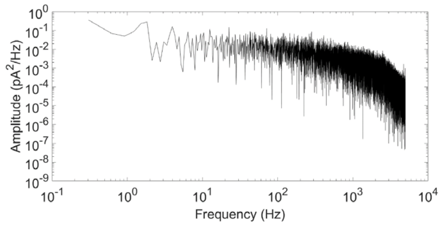

PBD22PEO14

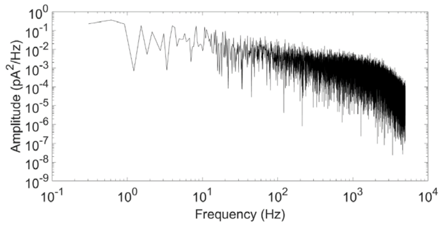

PBD11PEO8

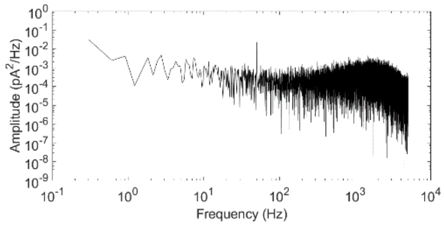

PBD11PEO8 + DPhPC (1:1 w/w)

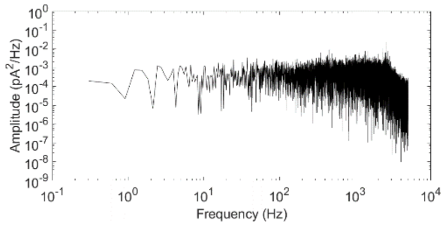

PBD22PEO14 + DPhPC (2:1 w/w)

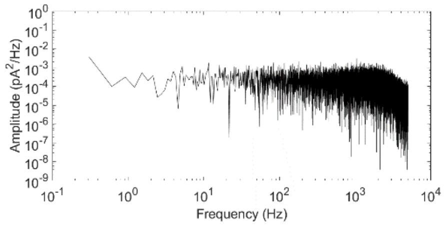

Figure S3: **Noise spectra of α-Hemolysin in DPhPC and all tested polymer-based membranes.** Noise spectra were obtained from electrical recordings at 75 mV, in a buffer with 1 M KCl, 15 mM TRIS and pH 7.5. Measurement was performed with 10 kHz sample frequency with a 2 kHz Bessel low-pass filter.

DPhPC

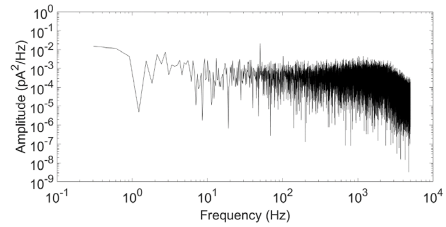

PMOXA$_4$PDMS$_{22}$PMOXA$_4$

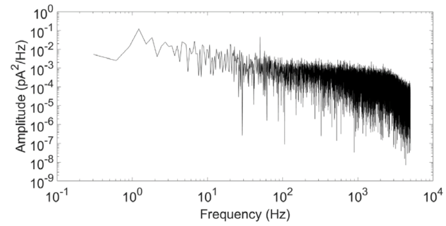

PMOXA$_{18}$PDMS$_{68}$PMOXA$_{18}$

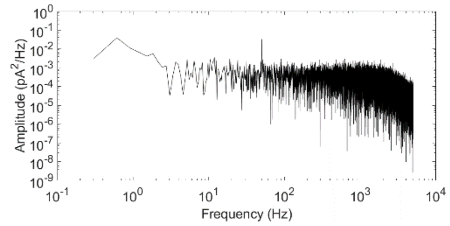

PBD$_{11}$PEO$_8$

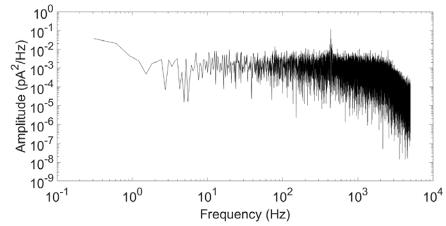

PBD$_{11}$PEO$_8$ + DPhPC (1:1 w/w)

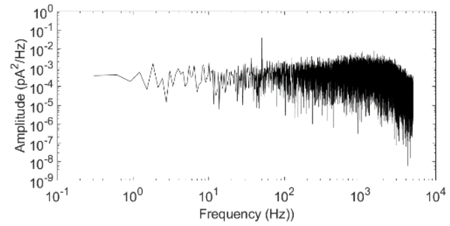

PBD$_{22}$PEO$_{14}$ + DPhPC (2:1 w/w)

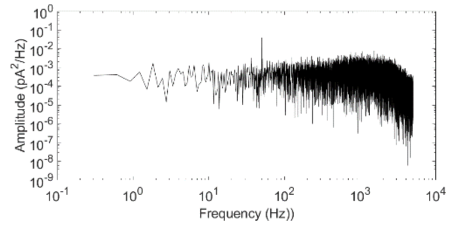

Figure S4: **Noise spectra of lysenin in DPhPC and all tested polymer-based membranes.** Noise spectra were obtained from electrical recordings at 75 mV, in a buffer with 1 M KCl, 15 mM TRIS and pH 7.5. Measurement was performed with 10 kHz sample frequency with a 2 kHz Bessel low-pass filter.

DPhPC

$PMOXA_4PDMS_{22}PMOXA_4$

$PMOXA_{18}PDMS_{68}PMOXA_{18}$

$PBD_{11}PEO_8$

$PBD_{11}PEO_8$ + DPhPC (1:1 w/w)

$PBD_{22}PEO_{14}$ + DPhPC (2:1 w/w)

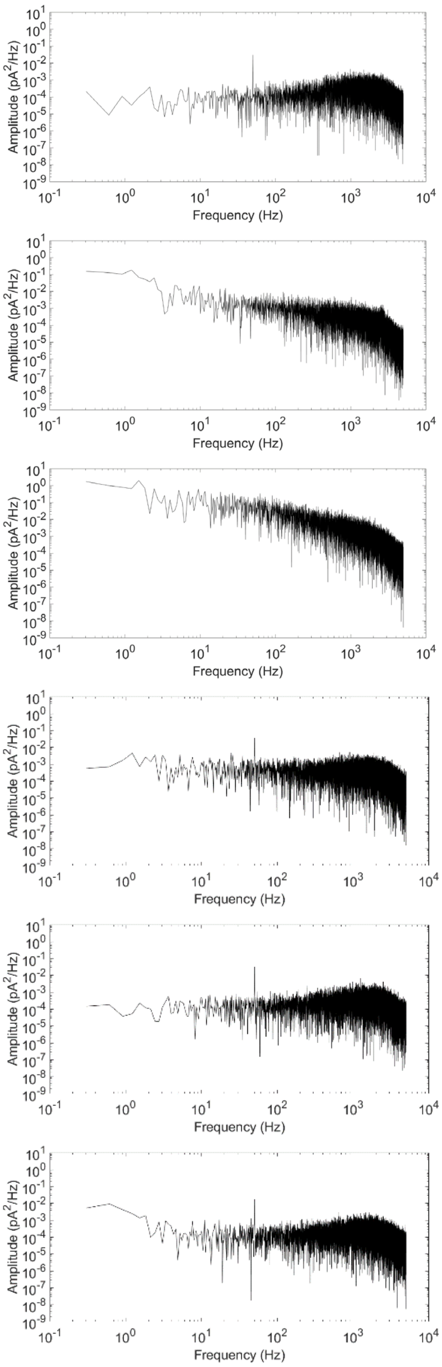

Figure S5: **Noise spectra of MspA-M2 in DPhPC and all tested polymer-based membranes.** Noise spectra were obtained from electrical recordings at 75 mV, in a buffer with 1 M KCl, 15 mM TRIS and pH 7.5. Measurement was performed with 10 kHz sample frequency with a 2 kHz Bessel low-pass filter.

DPhPC

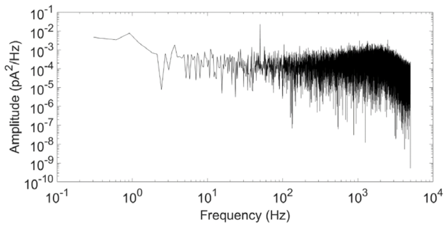

PBD22PEO14

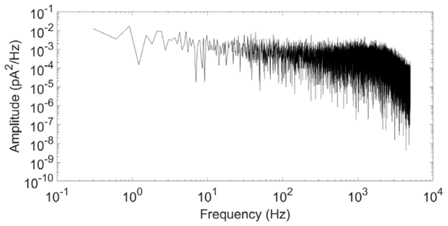

PBD11PEO8

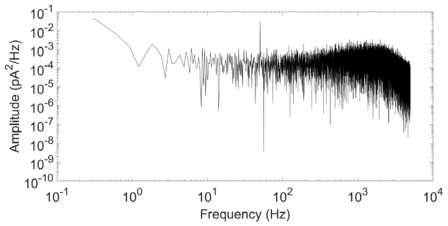

PBD11PEO8 + DPhPC (1:1 w/w)

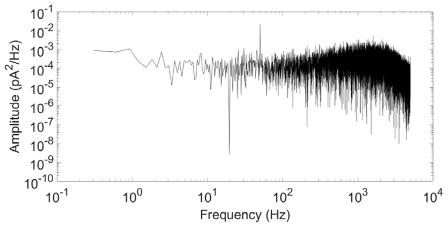

PBD22PEO14 + DPhPC (2:1 w/w)

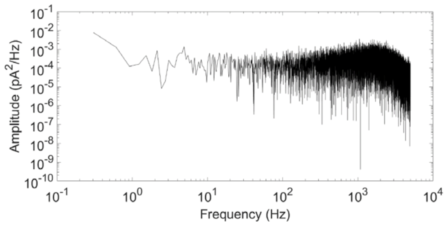

Figure S6: **Noise spectra of CytK in DPhPC and all tested polymer-based membranes.** Noise spectra were obtained from electrical recordings at 75 mV, in a buffer with 1 M KCl, 15 mM TRIS and pH 7.5. Measurement was performed with 10 kHz sample frequency with a 2 kHz Bessel low-pass filter.

DPhPC

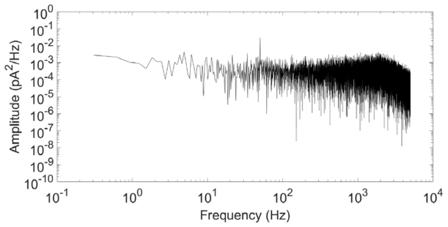

PBD22PEO14

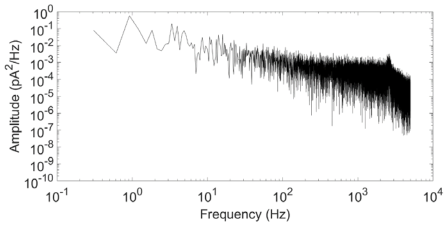

PBD11PEO8

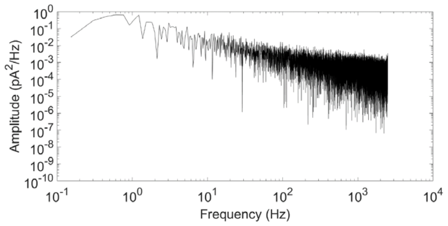

PBD11PEO8 + DPhPC (1:1 w/w)

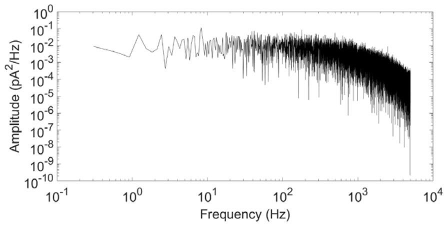

PBD22PEO14 + DPhPC (2:1 w/w)

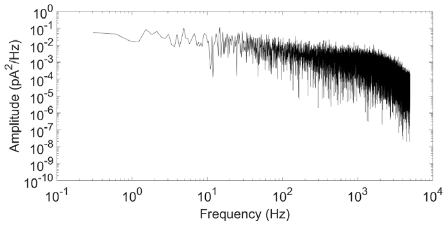

Figure S7: **Noise spectra of ClyA in DPhPC and all tested polymer-based membranes.** Noise spectra were obtained from electrical recordings at 75 mV, in a buffer with 1 M KCl, 15 mM TRIS and pH 7.5. Measurement was performed with 10 kHz sample frequency with a 2 kHz Bessel low-pass filter.

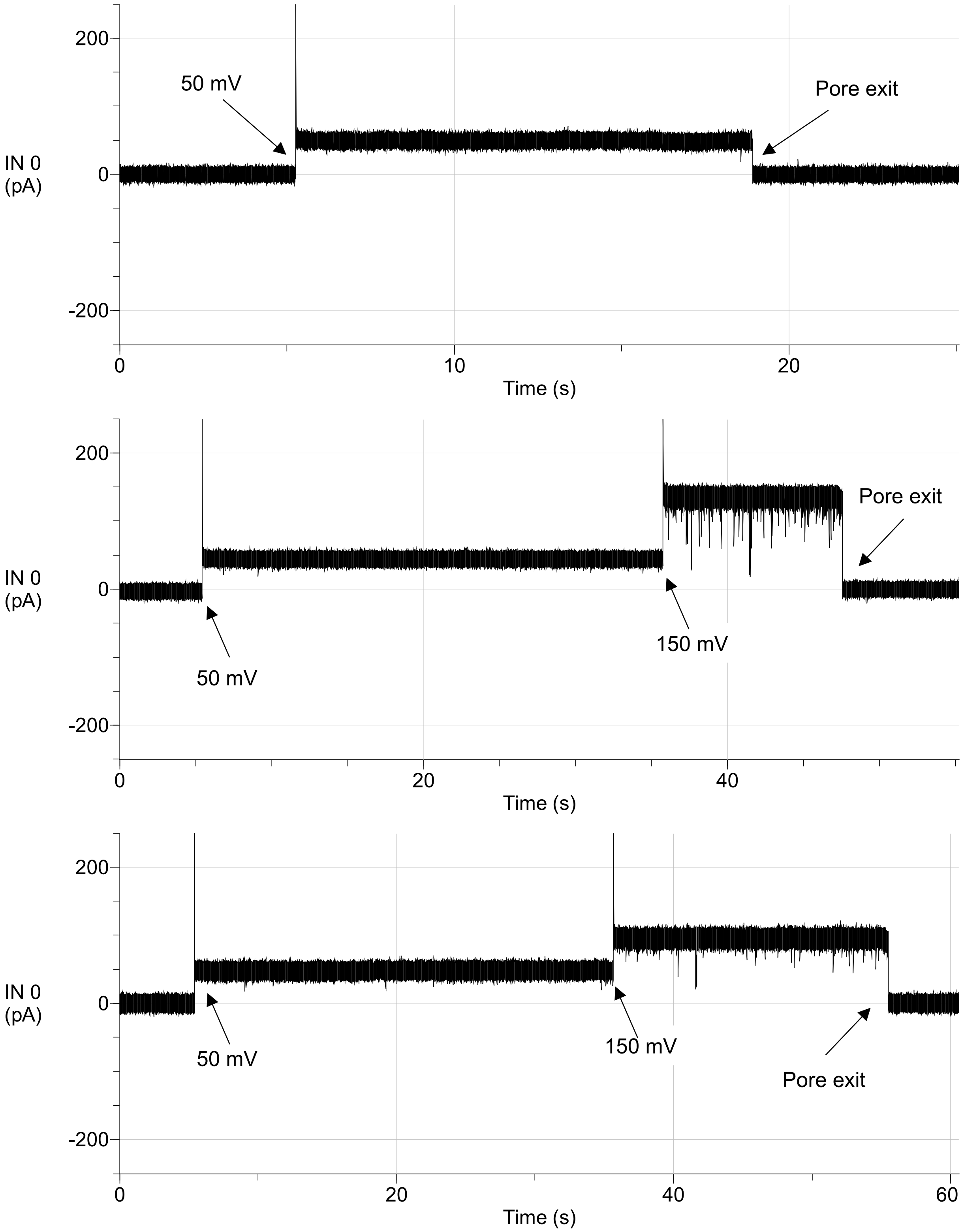

Figure S8: **Recording of α-hemolysin (top), aerolysin (middle) and MspA-M2 (bottom) exiting the $PBD_{22}PEO_{14}$ membrane shortly after the measurement was started.** Buffer conditions: 1 M KCl, 15 mM TRIS, pH 7.5.

DPhPC

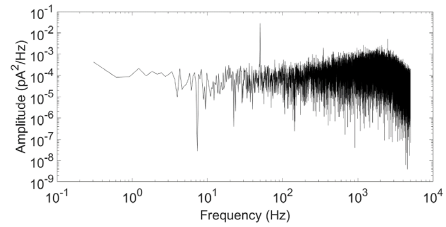

PBD$_{11}$PEO$_8$

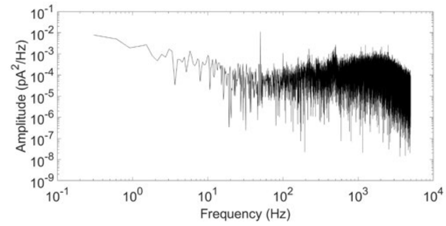

PBD$_{11}$PEO$_8$ + DPhPC (1:1 w/w)

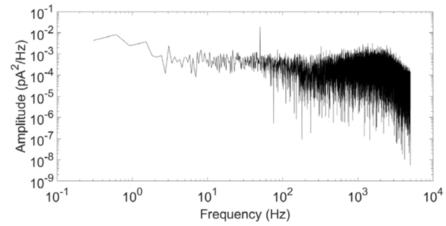

PBD$_{22}$PEO$_{14}$ + DPhPC (2:1 w/w)

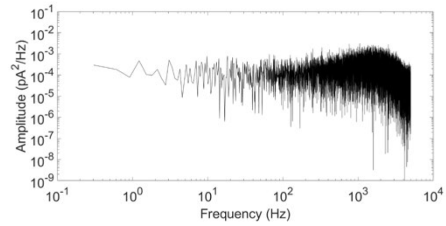

Figure S9: **Noise spectra of aerolysin in DPhPC and all tested polymer-based membranes.** Noise spectra were obtained from electrical recordings at 75 mV, in a buffer with 1 M KCl, 15 mM TRIS and pH 7.5. Measurement was performed with 10 kHz sample frequency with a 2 kHz Bessel low-pass filter.

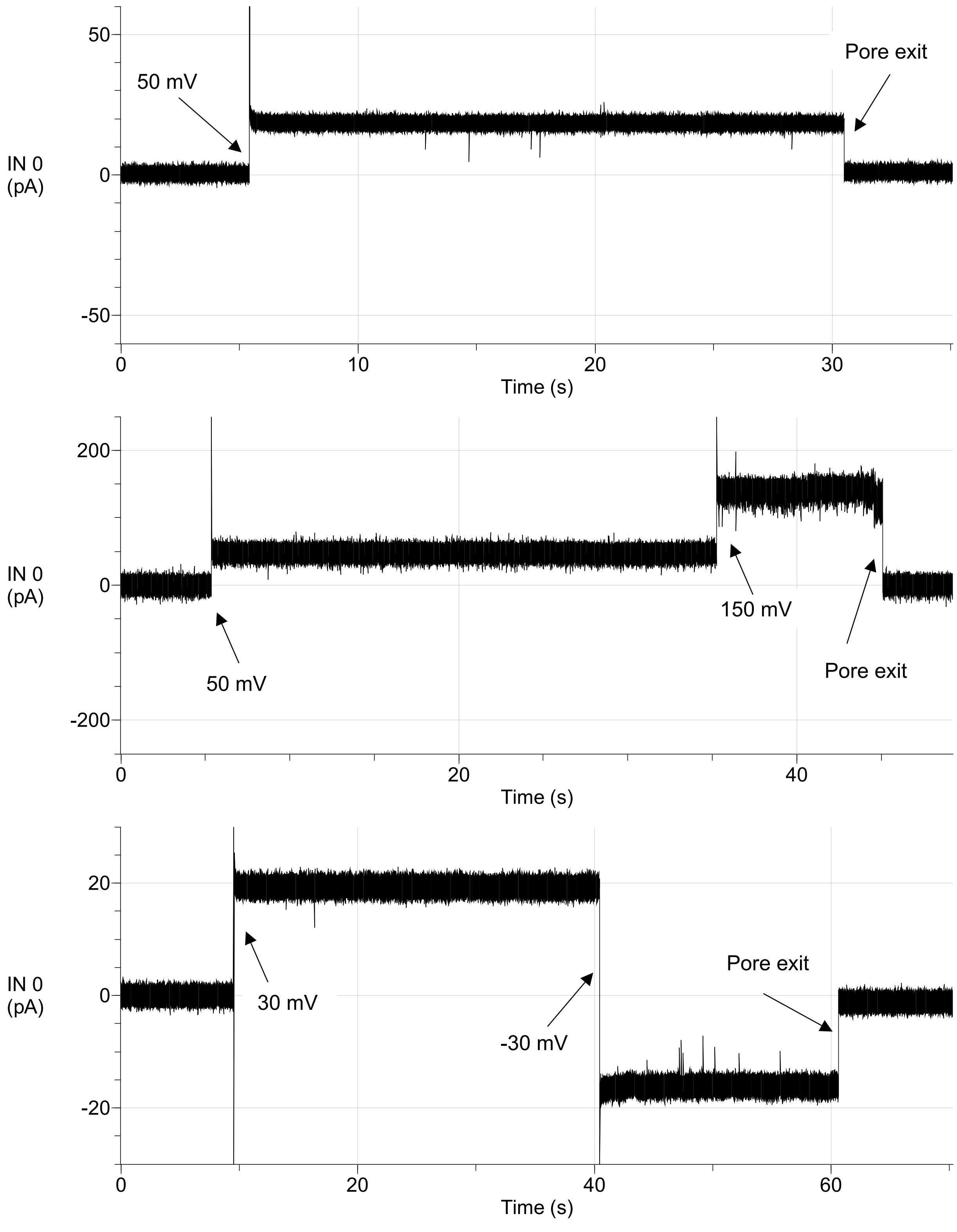

Figure S10: **Recordings of α-hemolysin (top), CytK (middle), and aerolysin (bottom), exiting the $PBD_{11}PEO_8$ membrane shortly after the measurement was started.** Buffer conditions: 1 M KCl, 15 mM TRIS, pH 7.5.

DPhPC

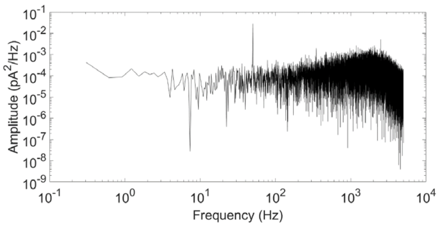

$PBD_{11}PEO_8$ + DPhPC (1:1 w/w)

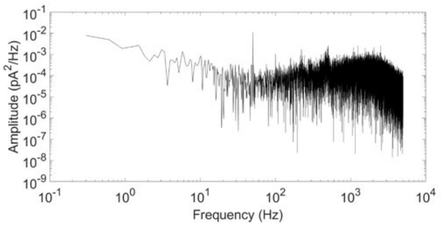

Figure S11: **Noise spectra of YaxAB in DPhPC and [$PBD_{11}PEO_8$ + DPhPC].** Noise spectra were obtained from electrical recordings at -75 mV, in a buffer with 150 mM NaCl, 50 mM TRIS and pH 7.5. Measurement was performed with 10 kHz sample frequency with a 2 kHz Bessel low-pass filter.

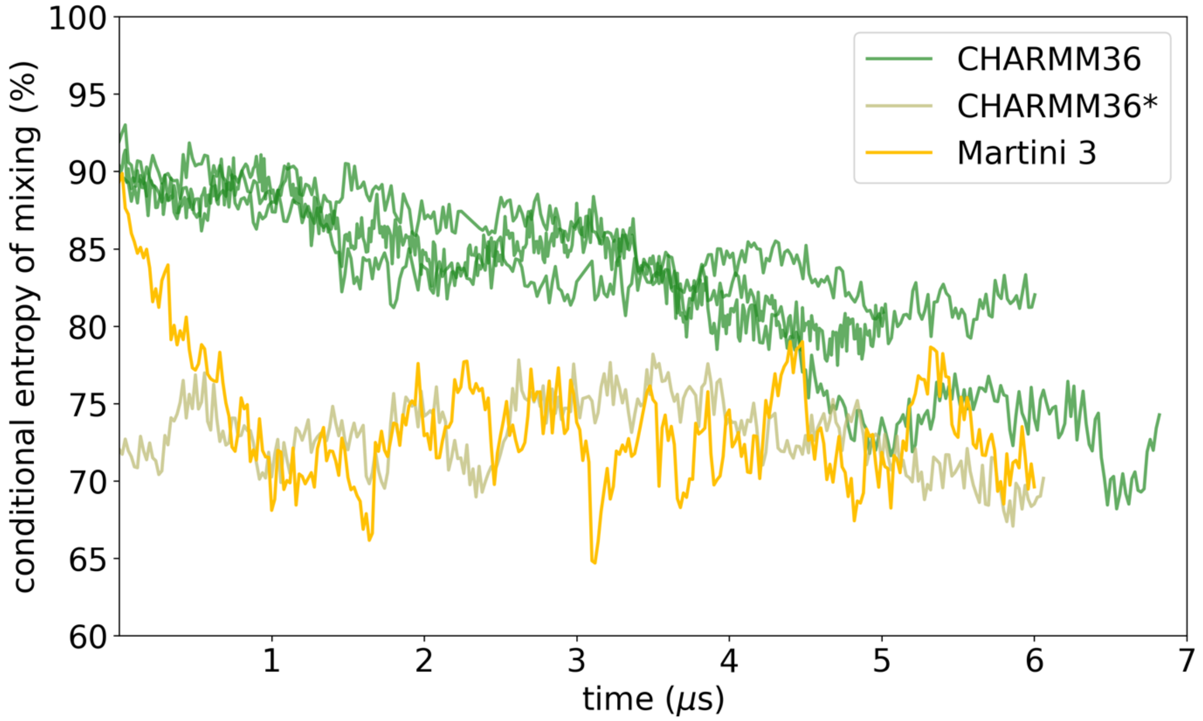

Figure S12: **Mixing of [$PBD_{11}PEO_8$ + DPhPC] hybrid membranes all-atom vs. coarse-grained.** Conditional entropy of mixing computed for four replicas of [$PBD_{11}PEO_8$ + DPhPC] using the CHARMM36 force field (green). One replica (*) was started by back mapping from an already converged Martini simulation. For reference a Martini 3 simulation with the same size (10 nm x 10 nm) is shown.

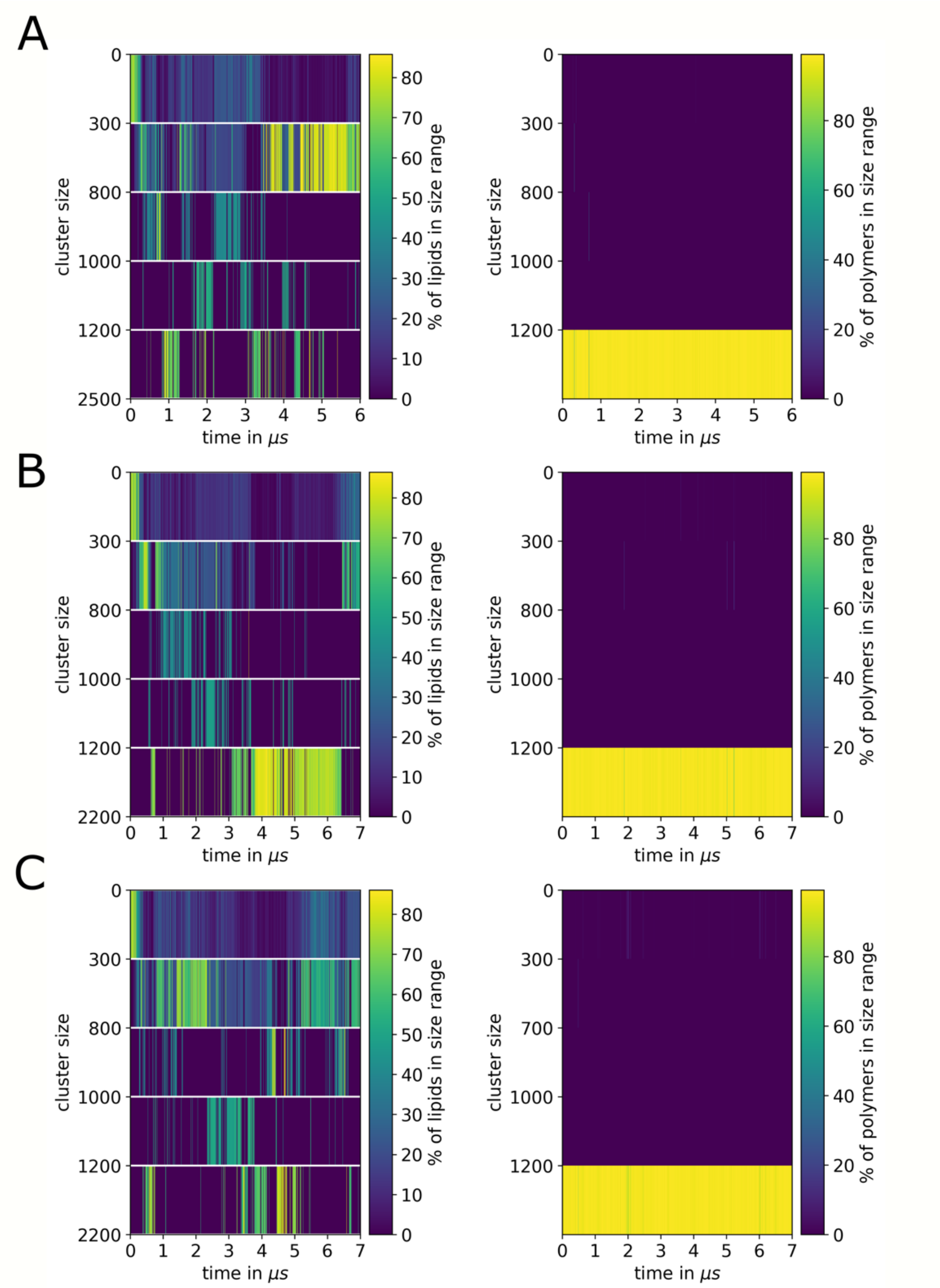

Figure S13: **Cluster analysis of 40 nm x 40 nm system.** The plots show the size ranges of clusters observed throughout the trajectory with the colour indicating how many percent of lipid or polymer molecules are located in clusters of a particular size range. For all replicas (A-C) the lipid clusters dynamically interchange between larger and smaller clusters. Replica A and C show an overall emergence of smaller clusters with sizes of 300-1000 lipids, while replica B shows a large cluster that only breaks apart towards the last microsecond. In contrast, the polymer molecules (right column) are forming one large connected cluster over the entire course of the simulation for all replicas.

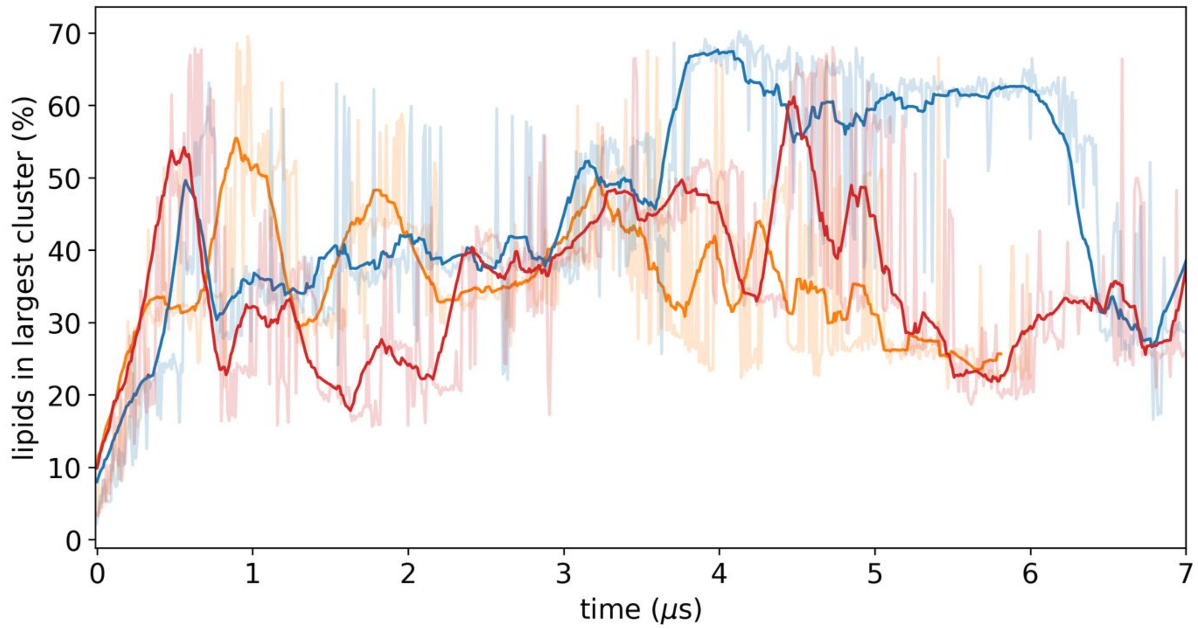

Figure S14. **Size of the largest cluster for three replicas (orange, blue, red) corresponding to A, B, C in Figure S13.** The running average (solid lines) shows that the first replica (orange) and the third replica (red) tend to have a maximum cluster, which contains 30-50% of all lipid molecules. In contrast, for the second replica the largest cluster contains around 60% of all lipid molecules starting at 3.5 µs which only breaks apart after 6.5 µs. Overall the maximum cluster size fluctuates dynamically indicating the interchange between larger and smaller nanodomains.

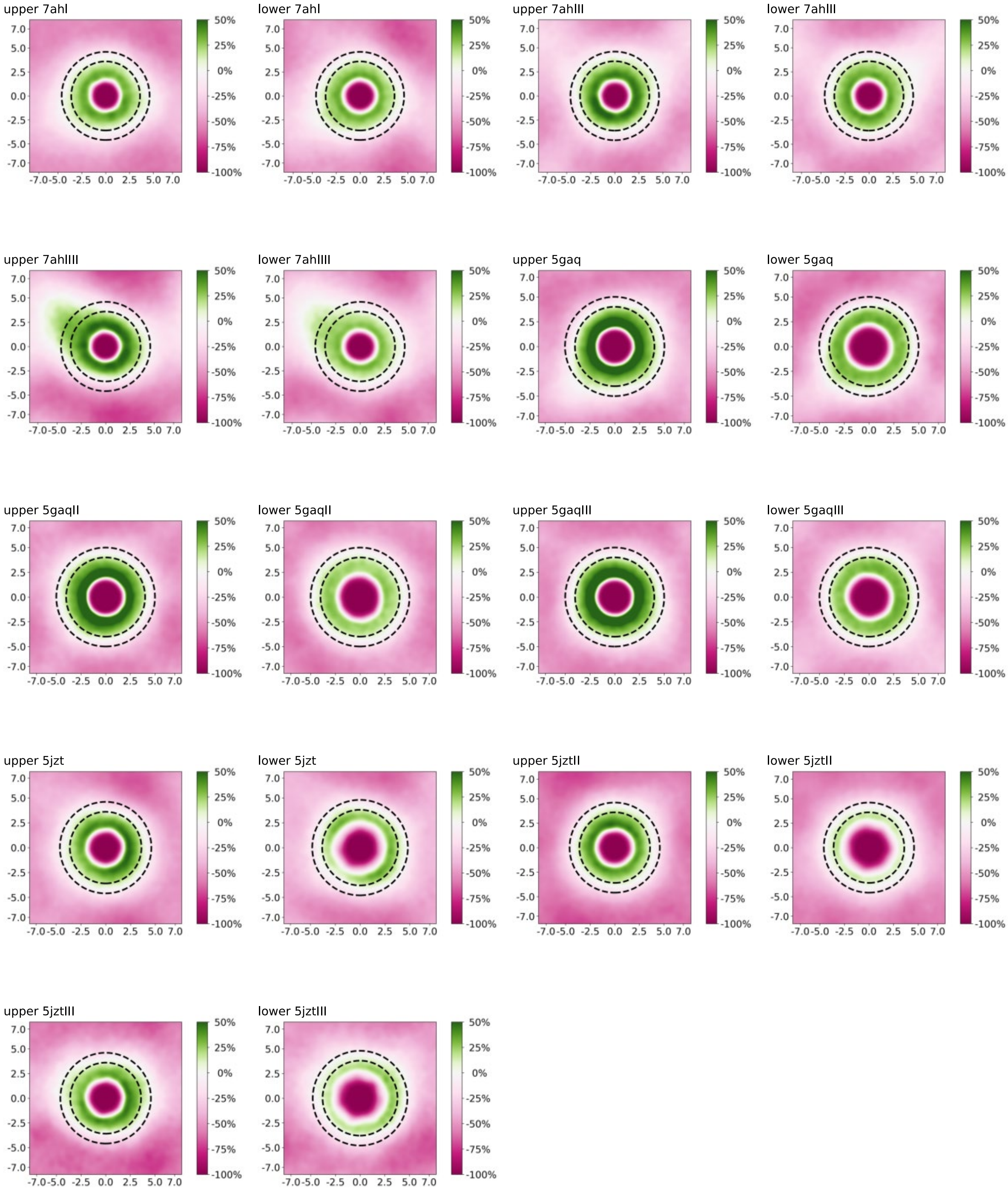

Figure S15. **Enrichment of DPhPC around proteins.** For each protein, three replicas were run and the enrichment was computed for the upper and lower leaflets. Green areas correspond to a positive enrichment (i.e. more lipids) and the pink areas to a depletion of lipids (i.e. less lipids). As expected, there is 100% depletion inside the pores as there are no lipids present in the channels.

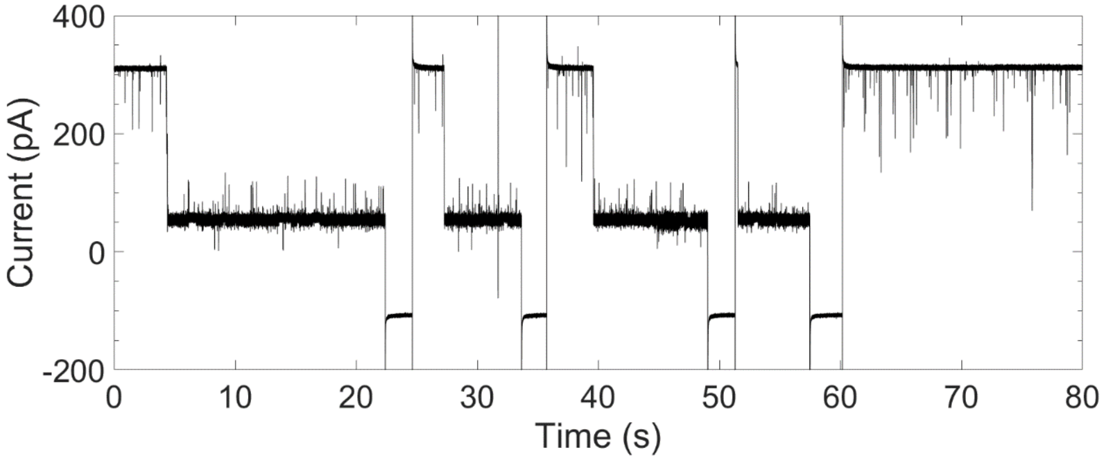

Figure S16: **Long-term blockade signals caused by [DNA-biotin:SA]-complexes with MspA-M2 in [$PBD_{11}PEO_8$ + DPhPC] hybrid membranes.** Fast blockades indicate the rapid translocation of 25-mer DNA-biotin across MspA-M2 nanopores. Moreover, long-lived blockade levels were observed, caused by DNA-biotin-SA complexes (25 nM added to *cis*). These complexes were unable to translocate, requiring the flipping of potential to -50 mV to remove the complex from the pore. Buffer conditions: 1 M KCl, 15 mM TRIS, pH 7.5. Recording was performed with 50 kHz sampling frequency, 10 kHz Bessel low-pass filter and for demonstration purposes the trace was additionally digitally filtered with a 2 kHz Gaussian filter.

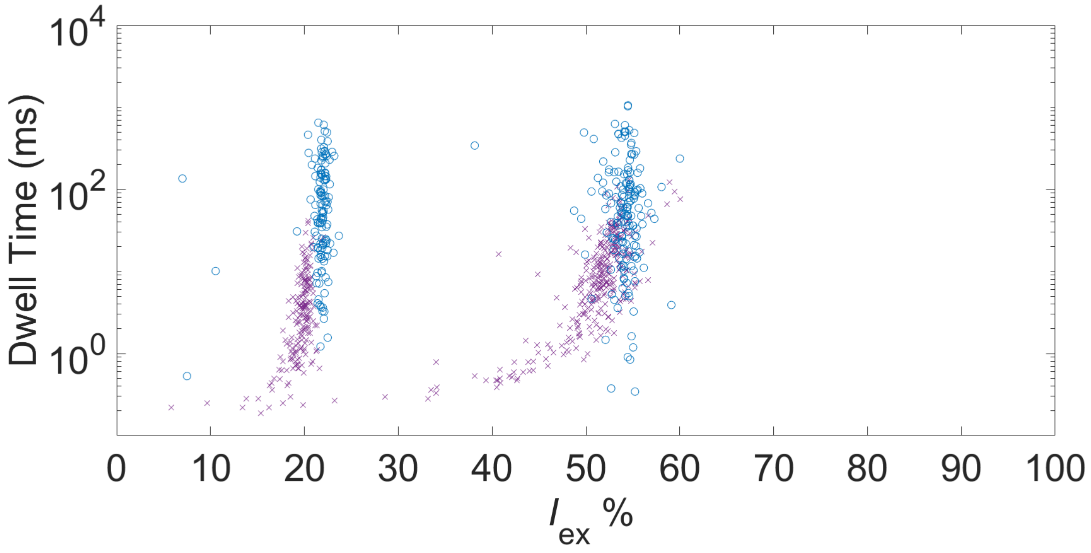

Figure S17: **Detection of SA and CRP with YaxAB in DPhPC and [$PBD_{11}PEO_8$ + DPhPC] hybrid membranes.** 0.25 µM of SA and 0.25 µM of CRP were added to *cis* and -75 mV was applied (150 mM NaCl, 15 mM TRIS, pH 7.5). Since SA (53 kDa) is able to penetrate deeper into the conically shaped pore than CRP (125 kDa), distinctive blockade signals are produced resulting in a higher excluded current ($I_{ex}$). When dwell time is plotted versus $I_{ex}$, two clearly separable clusters should arise that correspond to CRP (lower $I_{ex}$) and SA (higher $I_{ex}$), respectively. Blockades in DPhPC membranes (blue data points), were similar to blockades in [$PBD_{11}PEO_8$ + DPhPC] membranes (purple data points), showcasing that YaxAB maintains its conical pore shape in hybrid membranes, allowing the identification of differently sized proteins.

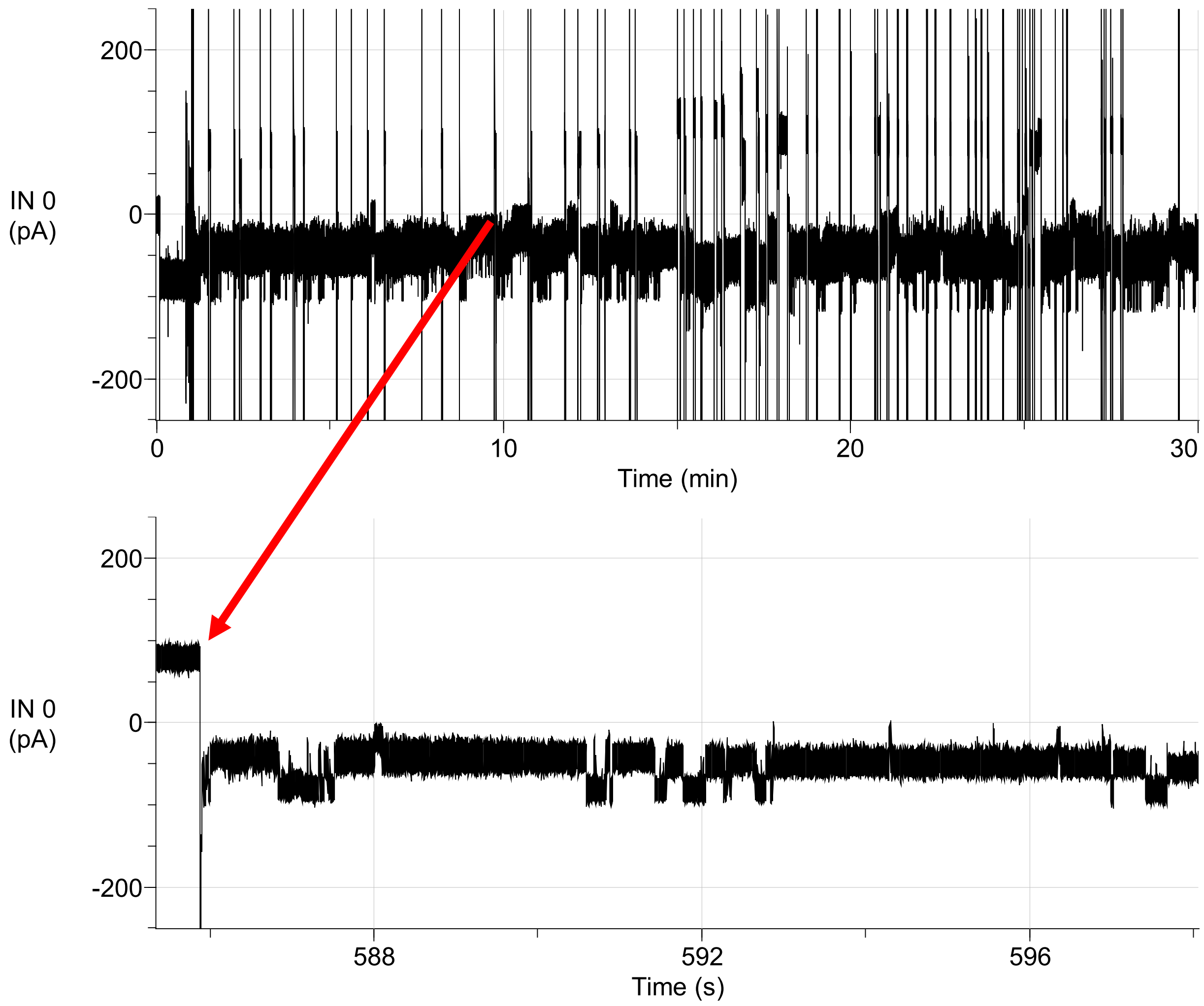

Figure S18: **30 min stable recording of YaxAB detecting 1:100 diluted human serum added to the *cis*-side at -50 mV.** The introduction of serum to the *cis*-chamber induces a high frequency of blockades. Long-lasting blocking events could be easily removed by briefly flipping the potential. Buffer conditions: 150 mM NaCl, 15 mM TRIS, pH 7.5. Recording was performed with 50 kHz sampling frequency, 10 kHz Bessel low-pass filter.

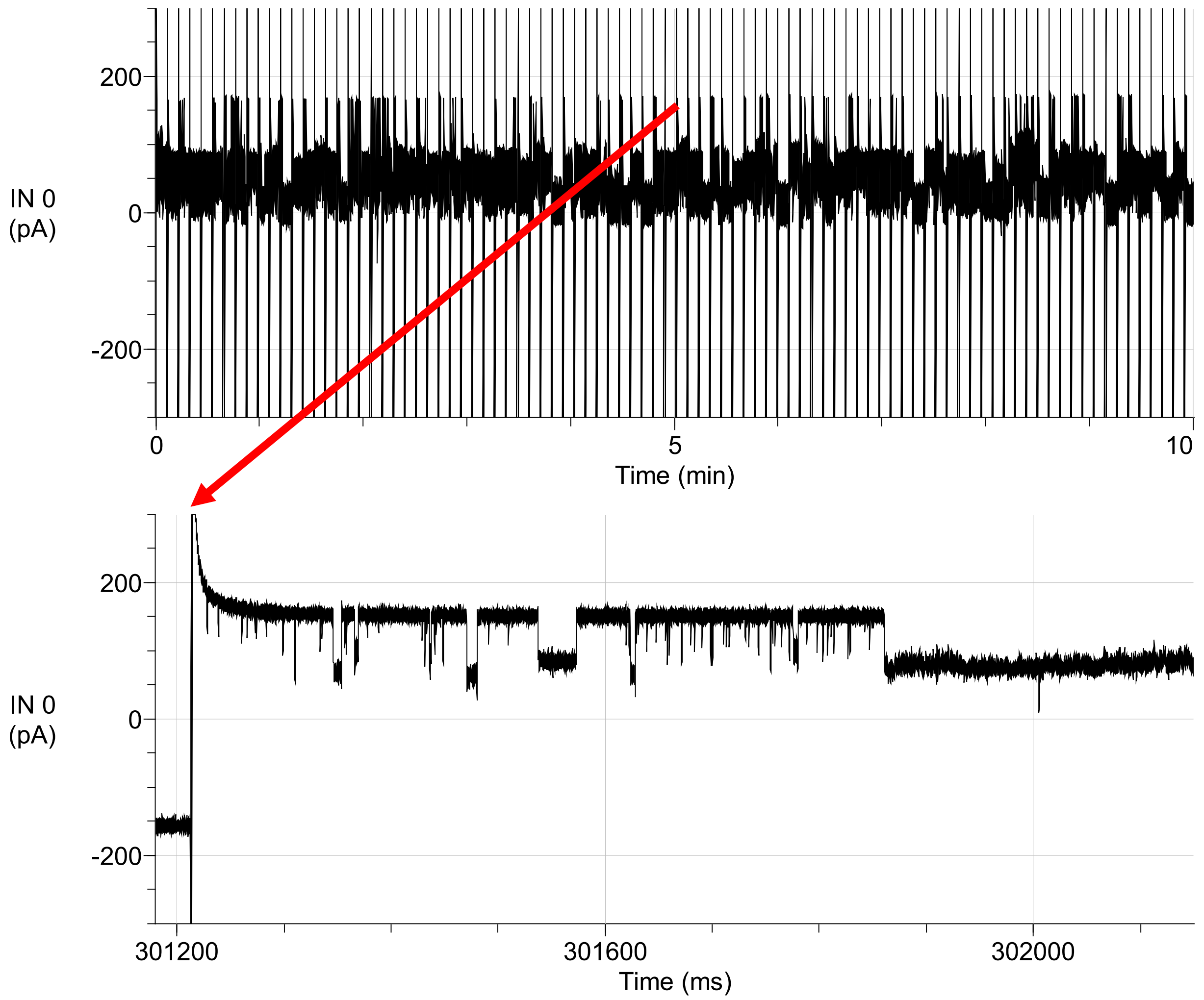

Figure S19: **10 min stable recording of lysenin detecting 25% v/v diluted human serum added to the *cis* and *trans* chambers at 75 mV.** The introduction of this high concentration of serum to lysenin created many blockades, caused by the many interactions of serum proteins with the pore. Long-lasting blocking events were often observed, but could be easily removed by briefly flipping the potential. Buffer conditions: 1 M KCl, 15 mM TRIS, pH 7.5. Recording was performed with 50 kHz sampling frequency, 10 kHz Bessel low-pass filter.
